# Functional and multiscale 3D structural investigation of brain tissue through correlative *in vivo* physiology, synchrotron micro-tomography and volume electron microscopy

**DOI:** 10.1101/2021.01.13.426503

**Authors:** Carles Bosch, Tobias Ackels, Alexandra Pacureanu, Yuxin Zhang, Christopher J Peddie, Manuel Berning, Norman Rzepka, Marie-Christine Zdora, Isabell Whiteley, Malte Storm, Anne Bonnin, Christoph Rau, Troy Margrie, Lucy Collinson, Andreas T Schaefer

**Affiliations:** Sensory Circuits and Neurotechnology Lab., The Francis Crick Institute, London, UK; Department of Neuroscience, Physiology and Pharmacology, University College London, UK; ESRF, The European Synchrotron, Grenoble, France; Electron Microscopy STP, The Francis Crick Institute, London, UK; Department of Connectomics, Max Planck Institute for Brain Research, Frankfurt am Main, Germany; scalable minds GmbH, Potsdam, Germany; Department of Physics and Astronomy, University College London, London, UK; Diamond Light Source, Harwell Science and Innovation Campus, Didcot, UK; School of Physics and Astronomy, University of Southampton, Highfield Campus, Southampton, UK; Paul Scherrer Institut, Villigen, Switzerland; Sainsbury Wellcome Centre, University College London, London, UK

## Abstract

Attributing *in vivo* neurophysiology to the brains’ ultrastructure requires a large field of view containing contextual anatomy. Electron microscopy (EM) is the gold standard technique to identify ultrastructure, yet acquiring volumes containing full mammalian neural circuits is challenging and time consuming using EM. Here, we show that synchrotron X-ray computed tomography (SXRT) provides rapid imaging of EM-prepared tissue volumes of several cubic millimetres. Resolution was sufficient for distinguishing cell bodies as well as for tracing apical dendrites in olfactory bulb and hippocampus, for up to 350 μm. Correlating EM with SXRT allowed us to associate dendritic spines on pyramidal cell apical dendrites in the stratum radiatum to their corresponding soma locations. Superficial pyramidal neurons had larger spine apparatus density compared to deeper ones, implying differential synaptic plasticity for superficial and deeper cells. Finally, we show that X-ray tomography and volume EM can be reliably correlated to prior *in vivo* imaging. Thus, combining functional measurements with multiscale X-ray microscopy and volume EM establishes a correlative workflow that enables functional and structural investigation of subcellular features in the context of cellular morphologies, tissues and ultimately whole organs.

## Introduction

For many decades, electron microscopy (EM) has been the dominant technique giving access to ultrastructural information in tissues and cells^1^. In neuroscience in particular, EM is at present the only technique to densely identify synapses. It is, however, primarily applied in 2D and in general limited in its spatial extent. Until the advent of high-throughput volume EM techniques^2^, electron microscopy allowed for extraction of ultrastructural information only over length scales of ~100 μm. Moreover, for obtaining insights in three dimensions, it entailed laborious manual serial sectioning, limiting the accessible depth to a few micrometers^3^ (with the notable exception of the heroic efforts of White, Brenner and colleagues^4^ assembling the *C. elegans* connectome from serial section TEM (ssTEM) over the course of ~15 years). Thus, while providing the highest resolution ultrastructural information, for mammalian tissue it has been difficult to obtain context information, such as the identity of neurons linked by synapses in an EM micrograph.

Fundamentally, two ways have been devised to overcome these limitations: Firstly, the revival and recent advances in volume electron microscopy, such as automated ssTEM^5^, serial blockface EM (SBEM)^6,7^, or focussed ion beam scanning EM (FIBSEM)^8^, have directly extended the imaged volume to up to several hundred micrometres in 3 dimensions, sufficient to capture the entire nervous systems of invertebrates^9^ or small vertebrate samples such as larvae^10,11^. In mammals, however, these approaches are in general still limited to small parts of brain regions^12^. Notably, in mammals it has been difficult to capture complete morphologies of individual neurons. Newly developed approaches such as multibeam scanning EM (mSEM)^13^, combined with automated serial sectioning^5,14^ or possibly gas cluster ion beam sectioning^15^, might in the future at least partially overcome some of these constraints. Nevertheless, these techniques remain highly time consuming (~months to years of data acquisition and processing times), so far rarely exceed volumes of 0.005 mm^3^, are largely reserved to a few specialist labs, and – due to the limited penetration depth of electrons in tissue – involve physical sectioning steps and are thus destructive and highly error-prone. Secondly, correlative methods can also provide context by linking local ultrastructural information to e.g. prior fluorescence measurements or by selectively labelling individual neurons using EM-compatible staining techniques^16^. Such approaches can give orthogonal information of functional/temporal (e.g. functional imaging prior to EM^17^) or molecular^18^ nature. They are, however, in general restricted to labelling and interrogating sparse subsets of cells. DNA barcoding-based techniques^19^ or super-resolution optical imaging^20^, possibly in combination with expansion microscopy^21^, might in the future enable at least somewhat denser segmentation of neurons and connections between neurons. Further challenges remain to link very different modalities with very distinct contrast mechanisms in efficient correlative pipelines^22^.

X-ray computed tomography (CT), i.e. 3D volumetric X-ray imaging, is a technique that can be useful in providing contextual information over large volumes: A single conventional X-ray CT scan can cover larger volumes than most volume EM approaches^23^. Furthermore, X-ray imaging relies on contrast mechanisms that are often analogous to those relevant in EM (elements that provide strong absorption contrast in X-ray images generally result in strong back-scattering or absorption signals in EM^23^). This in turn makes sample preparation compatible with both techniques and simplifies correlating datasets obtained by these two modalities. While the spatial resolution of laboratory X-ray systems can reach sub-micrometre length scales^24^, the broad polychromatic spectra and low spatial coherence of most laboratory systems can lead to blurring of features and image artefacts and can significantly limit the achievable image contrast and density resolution. This reduces the ability to reliably segment micrometre-sized structures and is only rarely sufficient to identify individual cell bodies^23,24^.

Synchrotron X-ray sources provide the ability to rapidly measure large volumes of a sample or large numbers of samples with good statistics^25^. Moreover, the use of X-ray phase-contrast (PC) imaging methods such as free-space propagation PC^26^, grating interferometry^27^, or speckle-based PC^28^ allows for measuring unstained biomedical samples in near-native state. Staining can be applied to further improve contrast^29–32^, and coherent imaging techniques such as X-ray holography^33^ bring the spatial resolution for biological tissue to the sub-100 nm level.

Here we demonstrate that parallel-beam synchrotron X-ray computed tomography (SXRT) can be used to obtain histological information from samples prepared for subsequent electron microscopy over length scales of several millimetres. The partially coherent X-rays allow for the use of phase-contrast imaging approaches^34,35^ enabling a spatial resolution sufficient to densely resolve neurites. The use of hard X-rays (with an energy higher than approximately 10 keV) in turn makes it possible to use heavy metal-rich stains that provide sufficient contrast also for subsequently performed volume EM experiments. Finally, SXRT, which is capable of investigating samples as large as several mm^3^ at sub-micrometre spatial resolution, combined with intermediate benchtop microCT^23^, which provides larger volumes at lower spatial resolution, can be integrated in an efficient multi-scale pipeline bridging from *in vivo* functional imaging experiments down to targeted volume EM of selected regions in a large histologically resolved volume.

## Results

To evaluate the use of synchrotron X-ray tomography (SXRT) for imaging neural circuits, we use the mouse olfactory bulb (OB) as a model system^36^. The OB is a well-ordered layered structure **(Fig. 1a-c)** that contains information at different length scales: Each receptor neuron (ORN) in the nose expresses a single olfactory receptor, out of 1000s possible. ORNs stretch their axons into the OB where those expressing the same type of receptor converge into 2-3 glomeruli, which are ~100 μm round neuropil structures. Beneath the layer of glomeruli is a ~200 μm thick external plexiform layer (EPL), largely containing straight dendrites and few cell bodies, followed by a thin layer of large somata of projection neurons (mitral cell layer) and a densely populated granular inner layer. As the OB is furthermore a superficial structure, accessible by *in vivo* functional imaging and has a number of diverse, prominent synaptic structures in the different layers, it offers a convenient model system to assess the limits of different imaging techniques and their combination **(Fig. 1, 2)**.

**Figure 1.**
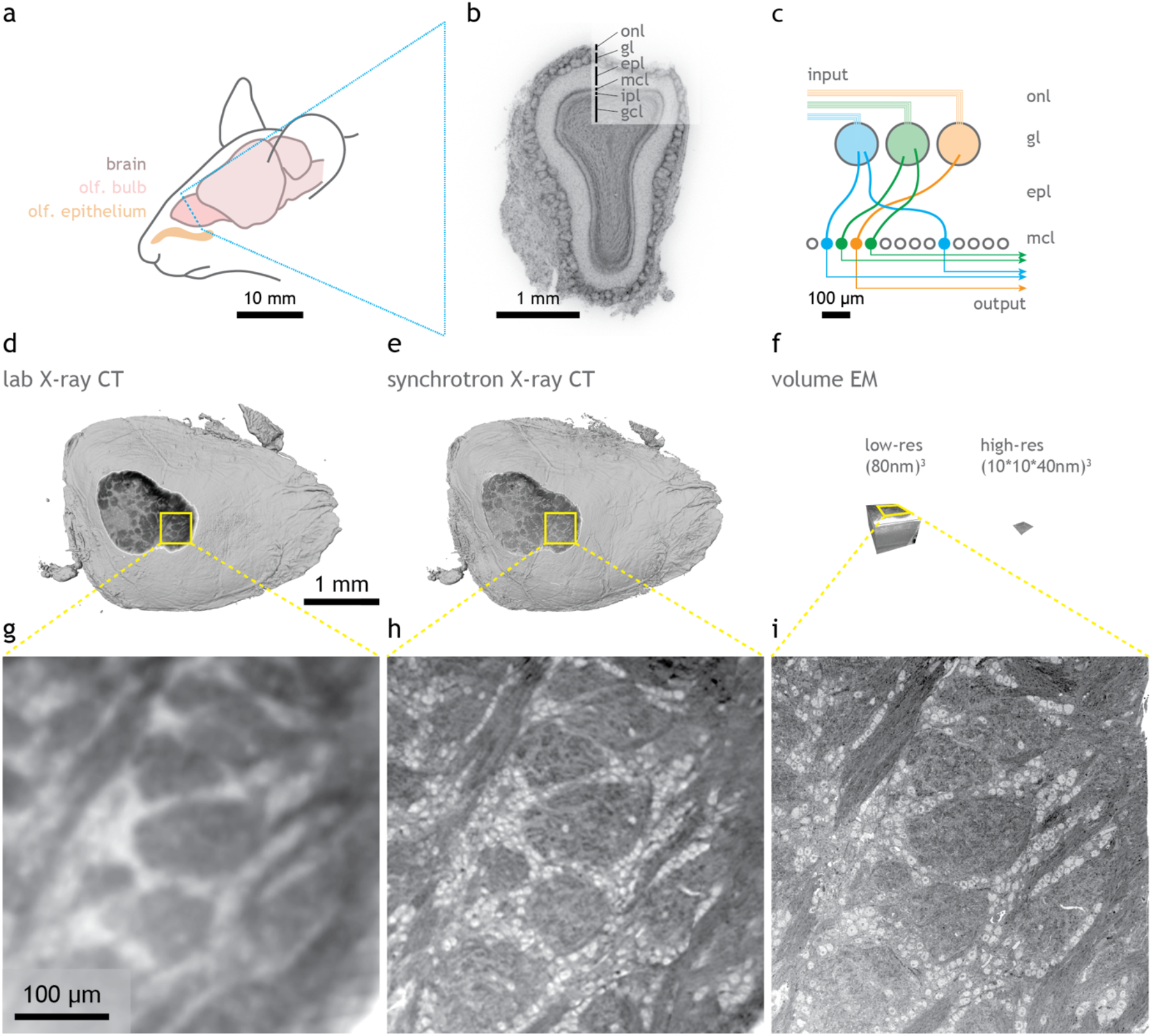
Correlative X-ray / volume electron microscopy for mouse brain tissue samples. **(a)** Elements of the olfactory sensory pathway in the mouse brain. **(b)** Nuclear staining of a 100 μm coronal section of the mouse main olfactory bulb, displaying its histological layers, obtained with widefield fluorescence microscopy. **(c)** Diagram of the excitatory wiring of the glomerular columns in the olfactory bulb. **(d-f)** Reconstruction at scale of the dataset volumes obtained by each imaging modality: lab X-ray CT **(d)**, synchrotron X-ray CT **(e)** and volume EM **(f)** at low ((80 nm)^3^) and high (10×10×40 nm^3^) resolution (left and right respectively), all warped to a common 3D framework. **(g-i)** Snapshots of the same region recorded by all three modalities. *onl*, olfactory nerve layer; *gl*, glomerular layer; *epl*, external plexiform layer; *mcl*, mitral cell layer; *ipl*, inner plexiform layer; *gcl*, granule cell layer.

**Figure 2.**
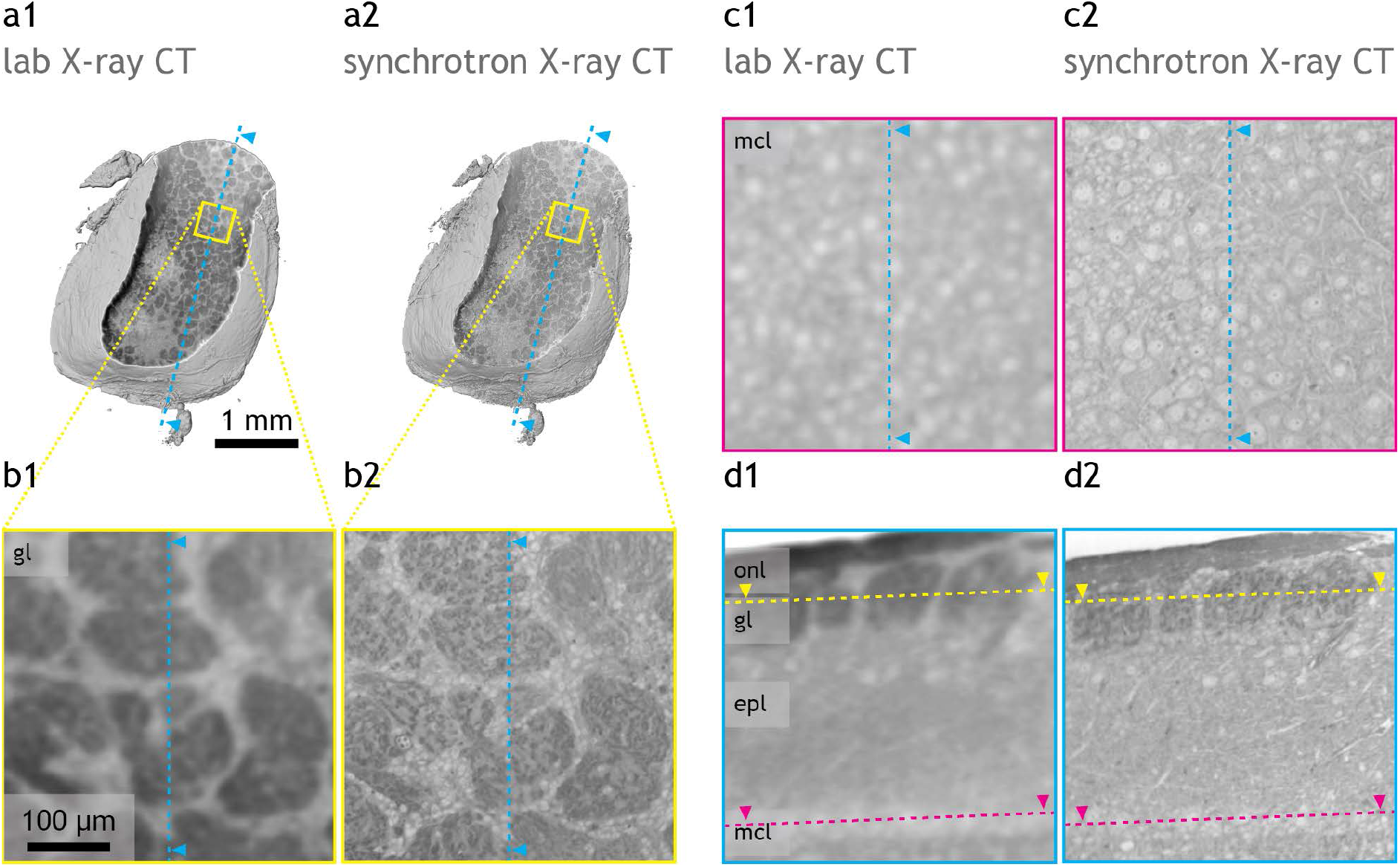
Lab and synchrotron X-ray CT of the same sample. **(a)** Reconstruction of the lab **(a1)** and synchrotron **(a2)** X-ray CT datasets obtained from the same sample, virtually sliced displaying the glomerular layer. **(b-d)** Virtual slices in three regions showing features resolved by each imaging modality in the glomerular layer **(b1-2)**, mitral cell layer **(c1-2)** and in a sagittal section **(d1-2)**. *onl*, olfactory nerve layer; *gl*, glomerular layer; *epl*, external plexiform layer; *mcl*, mitral cell layer.

We thus stained mouse olfactory bulb tissue sections with heavy metals using a protocol that preserves tissue ultrastructure^37^ in preparation for X-ray imaging **(Supp. Fig. 1).** As expected, a Zeiss Versa 510 benchtop micro-CT was readily capable of resolving individual glomeruli **(Fig. 2a1)**. While for example the thick layer of mitral cell somata could be identified with laboratory-based micro-CT imaging **(Fig. 2c1, d1)**, individual cell bodies were only rarely resolved **(Fig. 2c1)** and even the largest neurites remained undetected **(Fig. 2d1)**.

In order to assess the benefit of high-brilliance, partially coherent X-rays, we imaged the same samples using parallel-beam SXRT at the microtomography beamline I13-2 at the Diamond Light Source^38^**(Fig. 1e,h; Fig. 2a2-d2, Supp. Fig. 2)**. Comparison with laboratory-based micro-CT imaging revealed that SXRT provided a sufficient increase in spatial resolution to allow ready identification of different domains within each glomerulus **(Fig. 2b2)**. Furthermore, cell body identification of projection neurons was reliable **(Fig. 2c2,d2)**, and even small cell bodies (e.g. juxtaglomerular cell bodies in **Fig. 2b2**) were readily resolved. Moreover, apical dendrites of mitral cells were apparent in individual cross sections **(Fig. 2d2)**. To quantify to what extent SXRT is capable of delineating dendrites in general, we used tissue samples from both the hippocampus and olfactory bulb **(Fig. 3)**. After SXRT acquisition **(Fig. 3a1,a3,d1,d3)** we acquired serial block-face electron microscopy (SBEM) subvolumes at an isotropic voxel size of 50 nm (olfactory bulb) and at 80*80*50 nm^3^ (hippocampus) **(Fig. 3a4,d4)**. Both EM and SXRT datasets of each region were warped, allowing us to identify the same neuronal features in each modality **(Supp. Fig. 5)**. We then processed and imported both SXRT and SBEM datasets into the annotation tool webKnossos^39^ (see methods), where we seeded 50 mitral and 132 pyramidal cell somata and asked 3 tracers to independently trace apical dendrites as far as possible in both SXRT and EM data **(Fig. 3a2,b)**. In the OB, the majority of dendrites (29/44 traceable averaged across three tracers, see methods) were traced correctly (judged by comparison to the “gold standard” trace based on consensus SBEM data) and could be followed for up to 350 μm, often sufficient to traverse the whole EPL (mean traceable length averaged among 3 tracers = 153.6 μm) **(Fig. 3b,c)**. Re-imaging the same sample at a different SXRT beamline (TOMCAT at the Swiss Light Source) demonstrated that apical dendrite tracing was indeed readily reproducible (**Supp. Fig. 4**), suggesting that dendrite-level anatomical data can be efficiently achieved with many of the microtomography beamlines present at synchrotrons around the world.

**Figure 3.**
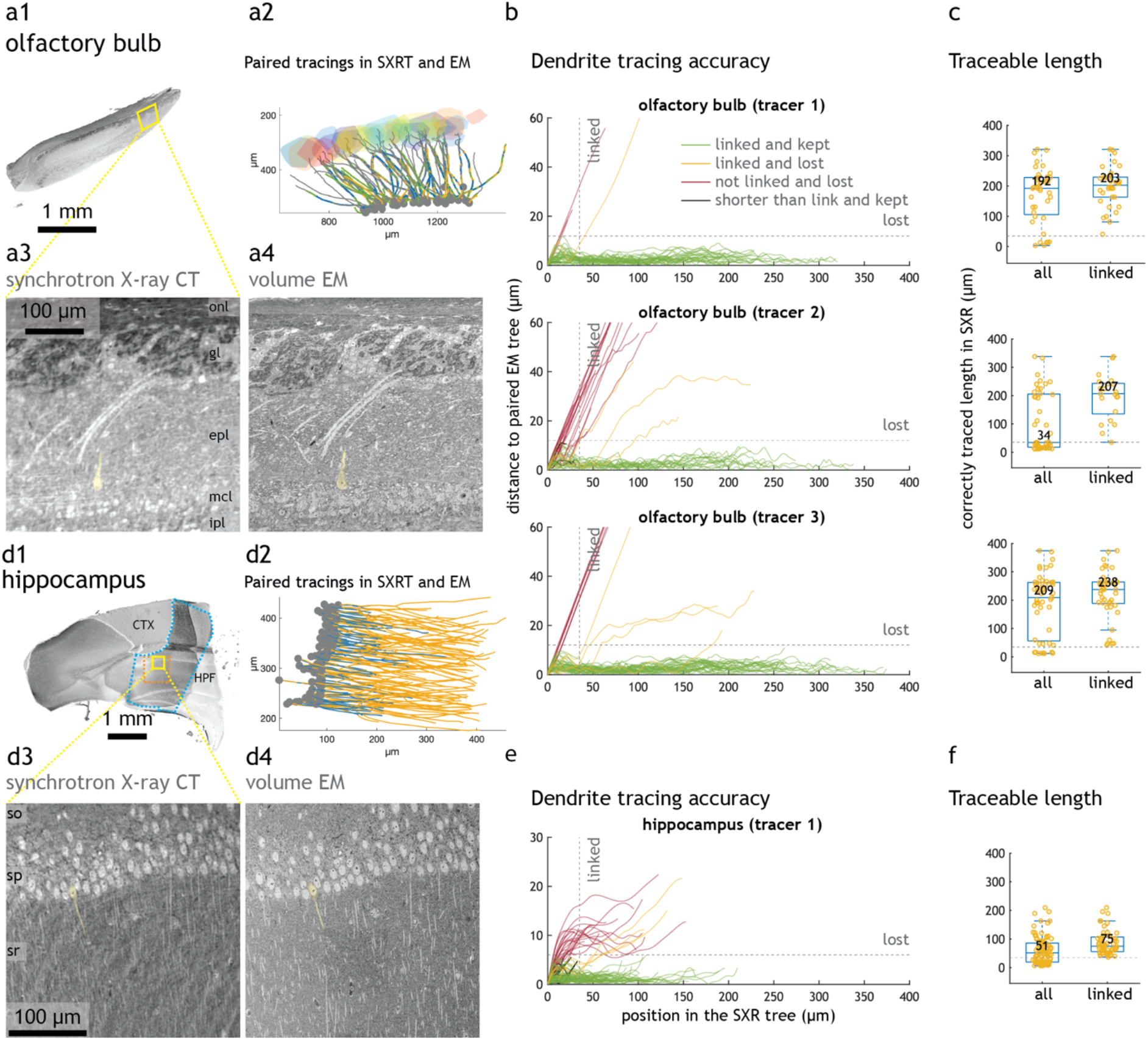
Apical dendrites of principal neurons can be traced in synchrotron X-ray CT datasets. **(a1)** Volume of a mouse brain olfactory bulb obtained with synchrotron X-ray CT, virtually sliced sagittally, displaying all main histological layers. Below, synchrotron X-ray CT **(a3)** and low-resolution volume EM ((50 nm)^3^ voxels) **(a4)** details of the highlighted region. Right, rendering of the same apical dendrites traced in both imaging modalities **(a2)**. **(b)** Distance between SXRT tracing and EM ground truth tracing of the same OB mitral cell apical dendrite evaluated at 320 nm steps along the entire traced dendrite. For each point on a SXRT dendrite tracing, the minimum distance to its paired EM tracing is plotted. (Prior to distance calculation SXRT and EM tracings were brought into the same dataset space, see methods). Each SXRT-EM pair of traces (i.e. one cell’s apical dendrite) is represented by a line, and is classified into four groups defined by two thresholds: a SXRT tracing is considered as “lost” if the average distance to its paired EM tree exceeds 12μm, otherwise as “kept”; a SXRT tracing is considered as “linked” if it is not lost before 35μm from the start of the tracing, otherwise as “not linked”. Lines belonging to each group are color-coded accordingly (green, yellow, red and black lines). Dashed lines indicate where linkage thresholds (35 μm) and lost thresholds (12 μm) are. Three tracers performed the same task, and their individual accuracies are plotted separately. **(c)** Traceable length of all SXRT dendrites and of the correctly linked ones. Traceable length measures for how long SXRT tracing is within 12um (lost threshold) away from paired EM tracing. Each dot represents one cell’s apical dendrite. The box itself covers the 25% to 75% percentile; the middle bar represents the median value (printed above); the whiskers extend to the most extreme datapoints that are not an outlier (defined as outside of 1.5x interquartile range). The grey dashed line marks the lost threshold used. **(d1)** Volume of a mouse hippocampus obtained with X-ray imaging. Region acquired with synchrotron X-ray CT is indicated with a blue dashed line. The region acquired with low-resolution volume EM (80×80×50 nm^3^ voxels) is indicated with an orange dashed line. Below, synchrotron X-ray CT **(d3)** and volume EM **(d4)** detail of the highlighted region. Right, rendering of the same apical dendrites traced in both imaging modalities **(d2)**. **(e)** Same as **(b)** for CA1 pyramidal neurons. Dashed lines indicate where linkage thresholds (35 μm) and lost thresholds (6 μm) are, to characterise individual dendrite tracings in terms of traced length and accuracy. Only one tracer performed the task. **(f)** Same as **(c)** for CA1 pyramidal neurons. *onl*, olfactory nerve layer; *gl*, glomerular layer; *epl*, external plexiform layer; *mcl*, mitral cell layer; *ipl*, inner plexiform layer; *so*, stratum oriens; *sp*, stratum pyramidale; *sr*, stratum radiatum.

Most mistakes (10/13, averaged across three tracers) happened directly at the linkage between soma and apical dendrite (red lines in **Fig. 3b**) where the contrast between intracellular space and extracellular space is weakest, due to higher prevalence of “darkly-stained” endoplasmic reticulum (**Supp. Fig. 3**). Moreover, in a few cases (2/44) the tracers stopped tracing before that linkage was solved (black lines in **Fig. 3b**). Excluding cells lost at this linkage point the expected traceable length became both larger and less variable (mean 204.9 μm, **Fig. 3c**). In the hippocampus, apical dendrites of pyramidal neurons could also be manually followed as they traversed the stratum radiatum, albeit with slightly shorter traceable length (mean 60.4 μm for all and 86.9 μm for linked cells, **Fig. 3e,f**), possibly due to the thinner diameter of those dendrites, the higher degree of similarity in the trajectories between dendrites compared to the OB, and variation in staining, amongst other factors.

While this indicates that, with appropriate contrast enhancing measures (e.g. staining), parallel beam SXRT can be used to delineate subcellular features (e.g. neurites) reliably for several hundreds of micrometres, high-resolution X-ray tomography in general is often limited by radiation damage to the samples^40–42^. To assess this, we performed high-resolution electron microscopy of the same samples after synchrotron X-ray imaging **(Fig. 4)**. Although the radiation dose applied during synchrotron tomography measurements was on the order of mega Grays, subsequent serial sectioning with a vibrating diamond knife^43^ and scanning EM (SEM) imaging were not noticeably impaired by pre-exposure to synchrotron X-rays **(Fig. 4c,d)**. Notably, as highly X-ray absorbing elements provided the strongest contrast in SEM imaging as well, contrast between the different imaging modalities was virtually identical **(Fig. 1,4)**, simplifying warping procedures **(Supp. Fig. 5)**. Direct comparison of volume EM and SXRT **(Fig. 4b,d)** demonstrated that the overall structure at the investigated length scale was unperturbed, and post-synchrotron sectioning and imaging was feasible with little to no detrimental effect of the X-ray exposure. To assess whether ultrastructure was maintained as well, we examined high-resolution EM images in selected regions of the olfactory bulb **(Fig. 4e,f)**. Fine axon bundles in the olfactory nerve layer were readily visible and visually unperturbed **(Fig. 4f2, Supp. Fig. 7)** as were cellular morphology and subcellular organelles in the glomerular layer **(Fig. 4f1-3)**. Finally, high-resolution EM in the EPL robustly showed apical dendrites **(Fig. 4f4-6)** and synaptic contacts **(Fig. 4f6)**, indicating that even extensive SXRT imaging for energies of 22 keV and doses in the range of 10^6^ Gy did not noticeably impact the ultrastructure at a level needed for e.g. connectomic analyses.

**Figure 4.**
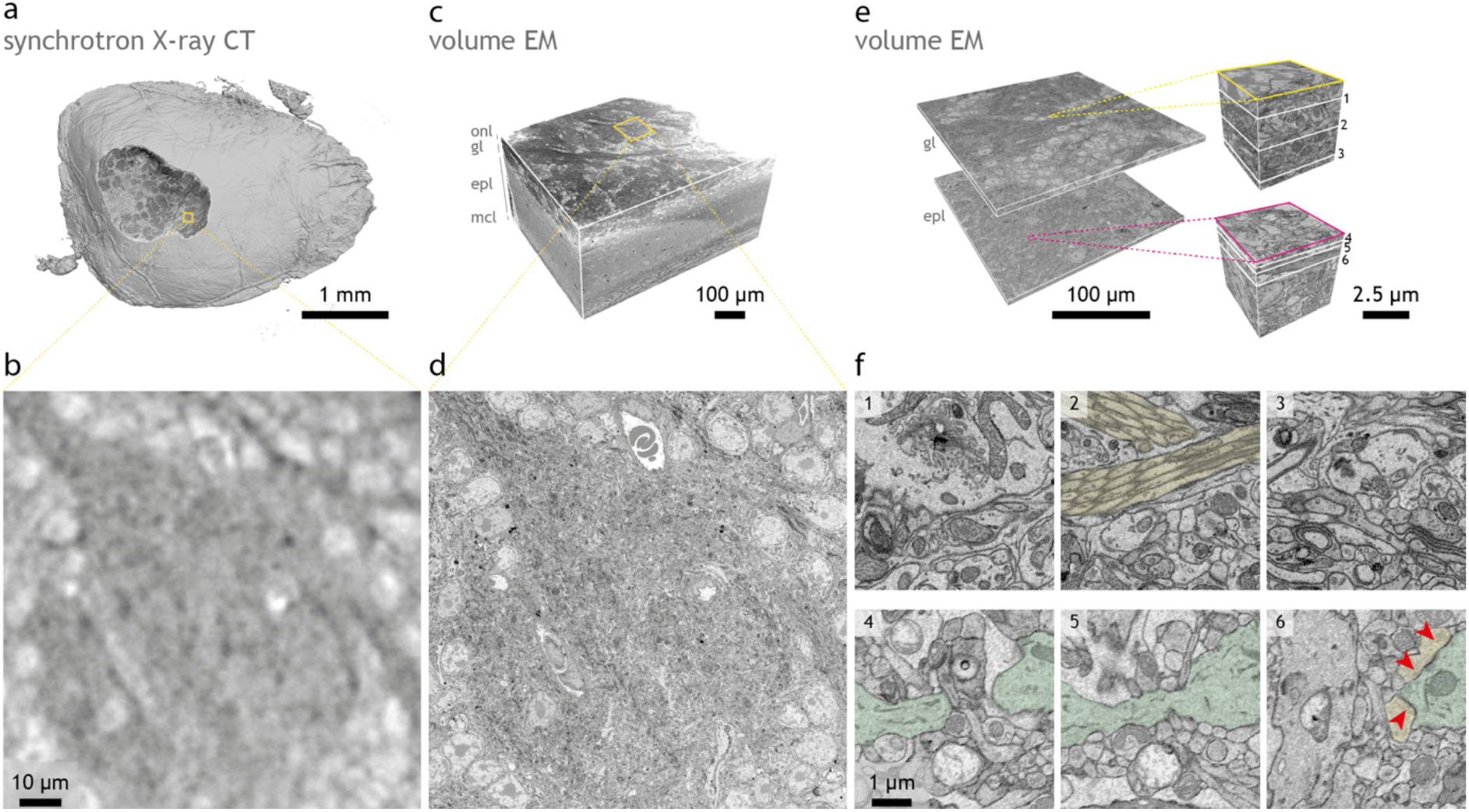
Tissue ultrastructure is preserved after synchrotron X-ray CT. Volumes obtained from the same sample: synchrotron X-ray CT **(a)** and posterior volume EM at low ((80 nm)^3^) **(c)** and high (10×10×40 nm^3^) resolution **(e)**, all warped to a common 3D framework. **(b,d)** Detail of the same region as detected by synchrotron X-ray CT **(b)** and by volume EM **(d)**. **(f)** Close-up details of the slices indicated in **(e)**, recorded by volume EM, displaying features indicative of well-preserved ultrastructure: axon bundles (yellow), thin dendrites (green) and synapses (red arrowheads). *onl*, olfactory nerve layer; *gl*, glomerular layer; *epl*, external plexiform layer; *mcl*, mitral cell layer

Together this suggests that SXRT has the potential to be used to provide histological, sub-cellular context to subsequent targeted, high-resolution EM imaging. To assess the feasibility of subsequent high-resolution EM imaging and correlation of the two datasets, we turned to the mouse hippocampus, a heavily-studied brain structure with clear layering and well-known synaptic circuitry. As expected, SXRT allowed us to identify pyramidal neuron cell bodies in the stratum pyramidale of CA1 (specifically subregion CA1a^44^, **Fig. 5a,b)**. As pyramidal cell apical dendrites could be readily traced through the stratum radiatum, we targeted the medial stratum radiatum with SBEM to analyse dendritic spines on selected dendrites **(Fig. 5a-c,g,j)**. We performed SBEM of a volume of 82 x 82 x 39 μm^3^ at a voxel size of 10 x 10 x 50 nm^3^. We then traced in the SBEM volume the apical dendritic tree of 7 pyramidal neurons previously identified in SXRT **(Fig. 5b,c)** and annotated a total of 3019 spines along these dendrites **(Fig. 5c)**. Among those 3019 spines, 13.8 % (417/3019) contained a spine apparatus **(Fig. 5j)**, a specialised endoplasmic reticulum organelle present in large spines and involved in calcium homeostasis and synaptic plasticity^1,45–48^. As suggested previously^49^, spines were denser on apical oblique dendrites compared to the trunk **(Fig. 5e)** as were spines with spine apparatus **(Fig. 5h)**. Relative prevalence was unchanged between trunk and oblique dendrites, and the distribution of both spines in general and spines with spine apparatus in particular – with few exceptions – largely followed a Poisson distribution **(Supp. Fig. 8)**.

**Figure 5.**
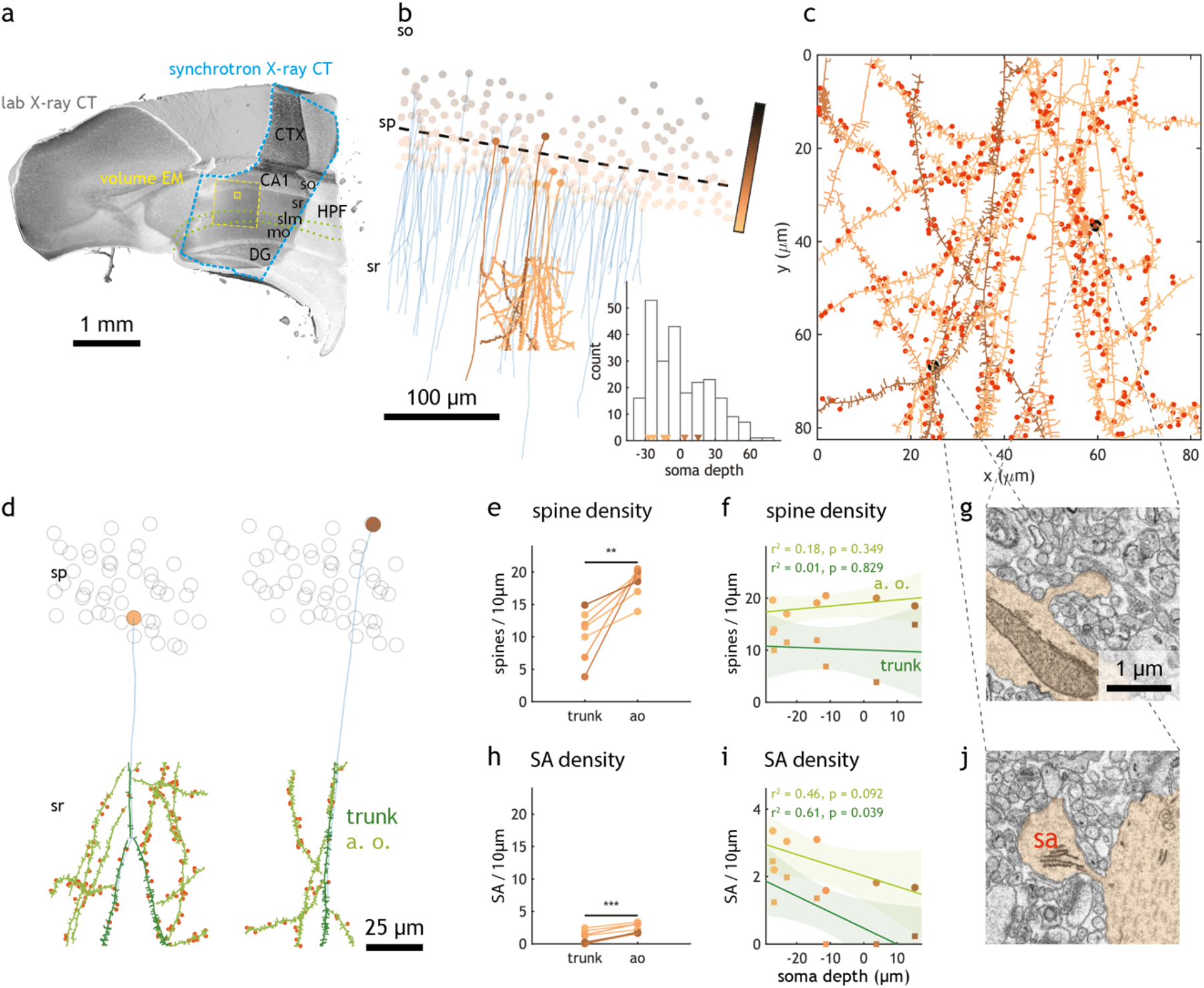
Multiscale analysis in the hippocampal CA1 region using correlative synchrotron X-ray CT / volume EM. **(a)** Reconstruction of a mouse hippocampus acquired with lab and synchrotron X-ray CT and volume EM (same as in Fig 3). The region acquired with synchrotron X-ray CT is indicated with a blue dashed line. The regions acquired with volume EM are indicated with yellow dashed (low resolution, 80×80×50 nm^3^ voxels) and yellow solid (high resolution, 10*10*50 nm^3^ voxels) lines. **(b)** CA1 pyramidal neurons, with their somata color-coded from brown to black according to soma depth in the pyramidal layer, and their apical dendrites traced in the SXRT dataset in blue. The dashed line shows a linear regression of the x,y coordinates of all seeded somata, which is used to quantify soma depth. The dendrites of seven cells whose apical dendrites entered the region imaged with high-resolution volume EM were traced exhaustively in EM and are shown in the same colour as their soma. The soma depths of all plotted somata are plotted as a histogram where the arrowheads point to the positions of the seven analysed neurons. **(c)** Dendrite traces of seven CA1 pyramidal neurons with spine-level details, segmented from the high-resolution EM dataset. All processes of the same colour belong to the same cell. Colour-coding represents soma depths as described in **(b)**. Spines are represented as little protrusions arising from the main skeleton which denotes the dendritic shaft. Spines with spine apparatus are highlighted in red. Two examples of spine and spine with spine apparatus are highlighted in black and their EM micrographs are depicted in **(g)** and **(j)**, respectively. **(d)** Examples of analysed CA1 pyramidal neurons. The apical dendrite traced in the SXRT dataset is shown in blue, dendrites traced in EM are shown in green (dark green: trunk; light green: apical oblique). Spines in these dendrites are shown as small protrusions, and those spines bearing spine apparatus contain a red dot. **(e)** Average spine density in trunk versus apical oblique dendrites (paired t-test), **(f)** plotted as a function of soma depth (colour coding as in **(b)**). The shaded area represents the 95% confidence interval of the linear regression line. **(h)** Average spine apparatus density in trunk versus apical oblique dendrites, **(i)** plotted as a function of soma depth, (* p<0.05, ** p<0.005, *** p<0.0005, ns non-significant in all panels). *CTX*, cerebral cortex; *CA1;* Ammon’s horn field CA1; *so*, stratum oriens; *sp*, stratum pyramidale; *sr*, stratum radiatum; *DG*, dentate gyrus.

While pyramidal neurons in the hippocampus are considered to be largely homogeneous, their properties vary along the spatial axes of this region. Differences affecting dendritic spines have been reported between pyramidal neurons with cell bodies located more superficial vs deeper in the pyramidal layer^44,50^. To assess whether dendritic spines in the stratum radiatum displayed different distribution or properties between neighbouring pyramidal cells, we linked the traces of apical dendrites in the high-resolution SBEM volume with those in the SXRT dataset, reaching the cell body layer **(Fig. 5b)**. This allowed us to give to each dendritic branch and spine their specific neural identity (i.e. more superficial or deeper). Spine density was independent of soma position **(Fig. 5f)**, but more superficial pyramidal cells had a substantially larger density of spines with spine apparatus **(Fig. 5i)**, possibly suggesting differential dynamics of calcium transients^45,47,48^. This analysis showcased that SXRT can indeed provide context – cellular identity – for EM annotations.

In order to corroborate that SXRT imaging can benefit the neuroscience community more broadly by providing subcellular context to subsequent high-resolution EM imaging, we performed SXRT imaging across a number of other brain regions, including the mouse neocortex, striatum, and cerebellum **(Fig. 6)**. Indeed, cell bodies, neurites and axon bundles were all readily visible in SXRT datasets of all brain regions, emphasizing the versatility of the approach.

**Figure 6.**
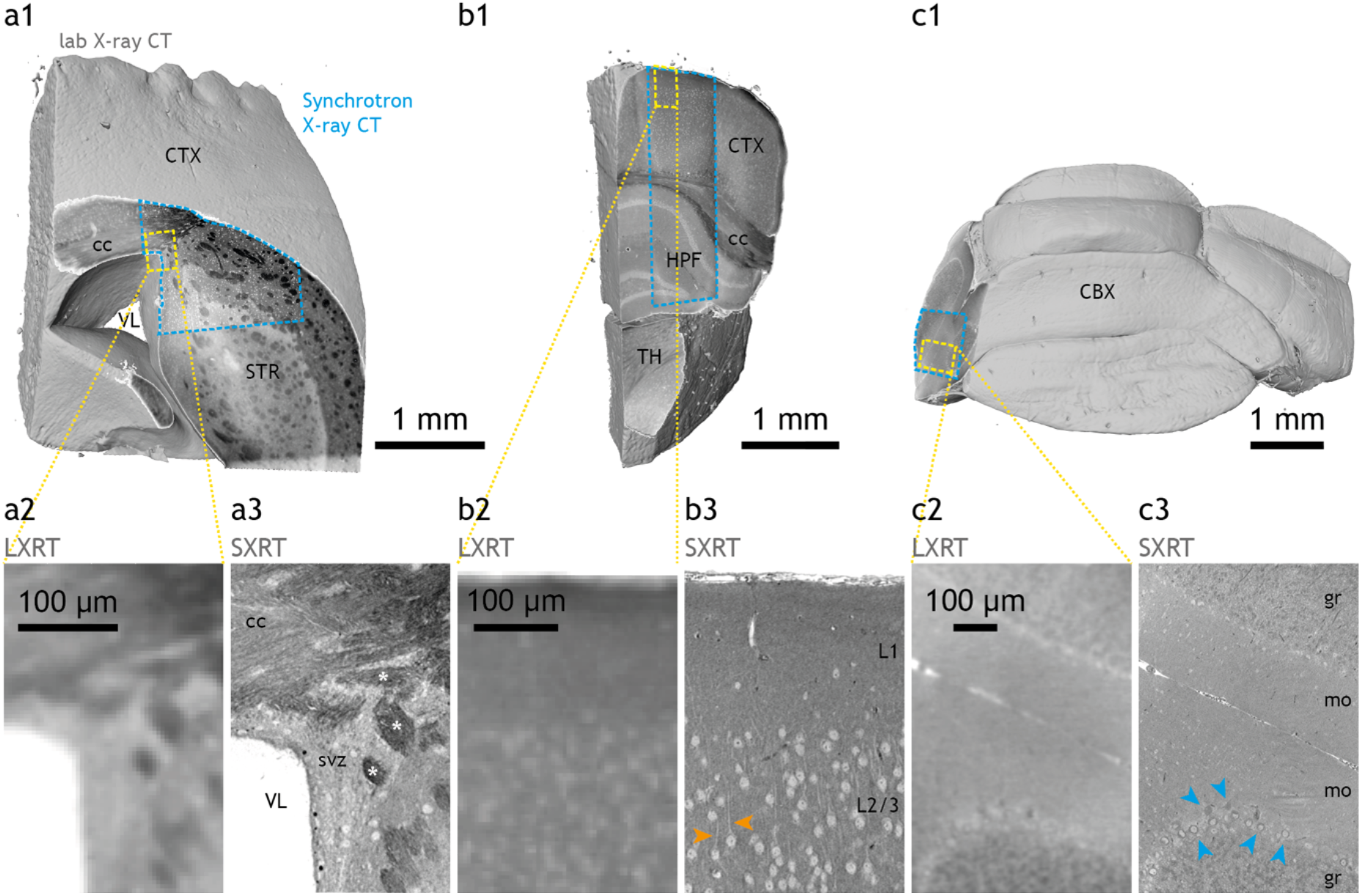
Subcellular compartments resolved by synchrotron X-ray CT in other mouse brain regions. Full-sample panels **(a1, b1, c1)** display synchrotron X-ray CT (SXRT) data where available (boundaries in blue dashed lines) and lab X-ray CT (LXRT) data elsewhere. Detail panels show lab **(a2, b2, c2)** and synchrotron X-ray CT **(a3, b3, c3)** data from the regions highlighted by yellow dashed lines in **(a1, b1, c1)**, respectively. **(a)** Mouse brain sample containing striatum, corpus callosum and cortex. Detail **(a3)** shows the subventricular zone, somata and myelinated axon bundles (white asterisks). **(b)** Sample containing cerebral cortex, hippocampus and thalamus. Detail **(b3)** shows cortical layers 1 and 2/3, displaying somata and apical dendrites of pyramidal neurons (orange arrowheads). **(c)** Sample containing cerebellar cortex. Detail **(c3)** shows a section across cerebellar lobules, displaying granular and molecular layers and purkinje neurons (blue arrowheads). *CTX*, cerebral cortex; *STR*, striatum (caudoputamen); *VL*, lateral ventricle; *cc*, corpus callosum; *svz*, subventricular zone; white asterisks, myelinated axon bundles; *HPF*, hippocampal formation; *TH*, thalamus; *L1*, cortex layer 1; *L2/3*, cortex layer 2/3; Orange arrowheads, apical dendrites of cortical pyramidal neurons; *CBX*, cerebellar cortex; *gr*, granular layer; *mo*, molecular layer; blue arrowheads, Purkinje neurons

One of the key challenges in mammalian circuit neuroscience is to bring together large-scale functional studies *in vivo* with their structural anatomical underpinning^17,51^. However, relating ~10-100 μm EM datasets to their corresponding ~mm scale *in vivo* imaging datasets is difficult: Full neural circuits are embedded in volumes often exceeding several mm^3^, making it challenging to acquire and analyse using EM^52^. Different length scales and staining protocols associated to the different imaging modalities of this multi-step process often result in structural distortions that need to be untangled to robustly correlate features across datasets. We hypothesised that the large volumes and high level of structural detail available with SXRT should substantially simplify this correlative pipeline.

To assess this and directly link *in vivo* functional data to the underlying anatomical features, we performed *in vivo* 2-photon Ca^2+^ imaging experiments in the mouse olfactory bulb. Presenting a panel of odours to anaesthetised mice^53^ revealed diverse response profiles of about 20 glomeruli on the dorsal OB surface **(Fig. 7a-e)**. Following *in vivo* functional imaging, we labelled blood vessels via an intraperitoneal injection of sulforhodamine (SR101) and performed anatomical fluorescence imaging of the entire tissue volume with ~0.36 μm horizontal and 5 μm vertical spatial resolution. Afterwards, mice were sacrificed, the OB extracted and a 600 μm dorsal section including all imaged glomeruli dissected. Autofluorescence after fixation allowed us to readily identify individual glomeruli and numerous landmarks that allowed for straightforward warping of the fluorescence dataset to datasets obtained by subsequent lab and synchrotron X-ray CT **(Supp. Fig. 5)**. This process is efficient enough to allow linking several features *in vivo* (e.g. different labelled glomeruli, **Fig. 7f-n**) to subsequent targeted EM **(Supp. Fig. 9)**. As before, laboratory- and synchrotron-based X-ray imaging and EM revealed different levels of detail of the glomerular anatomy with SXRT able to delineate essentially all glomeruli in the field of view (as gauged by EM acquisition of a subvolume, **Fig. 7m,n**). Relating glomeruli annotated in EM to the *in vivo* 2-photon imaging revealed that 3/20 glomeruli were misidentified *in vivo* with only X-ray or EM datasets allowing to correctly attribute fluorescence to anatomical units **(Fig. 7m,n, Supp. Fig. 10, Supp. Fig. 11)**.

**Figure 7.**
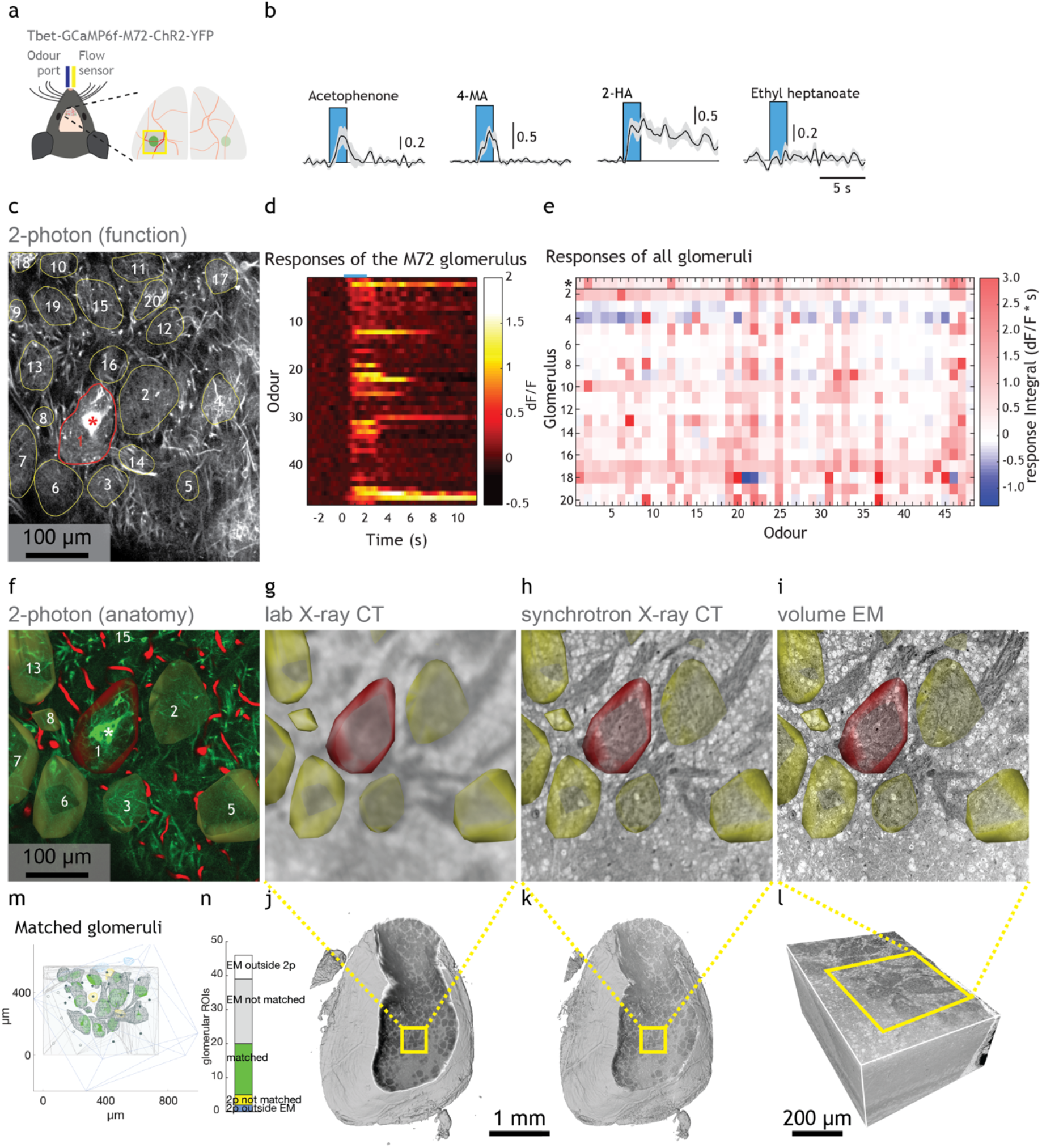
Multiscale analysis in the olfactory bulb glomerular layer using correlative in vivo 2-photon Ca^2+^ imaging / synchrotron X-ray CT / volume EM. **(a)** *in vivo* 2-photon imaging setup for imaging neural activity of projection neurons in anesthetised mice in response to a controlled odour pattern delivered to their nose. Right: Schematic of the olfactory bulb surface with blood vessels (red) and genetically targeted M72 glomerulus (green) indicated. **(b)** Example traces (mean of 5 presentations +/− SEM) of neuronal activity in response to four odours (odour presentation window in blue). **(c)** Maximum projection (8000 frames) of the functionally imaged plane recorded *in vivo,* located in the glomerular layer. The highly branched dendritic tufts allow delineating the contours of 20 putative glomeruli (yellow contour). The fluorescence signal inside these regions of interest was measured. In the M72 glomerulus the afferent sensory axons expressed YFP, making it identifiable (red asterisk and contour). **(d)** Neural activity in the M72 glomerulus in response to a panel of 48 odours (mean of 5 presentations +/− SEM). **(e)** Response integral of all glomeruli to all odours (average of 5 presentations). **(f)** Anatomical dataset obtained *in vivo* at the end of the functional recordings using 2-photon microscopy. (green: GCaMP / M72-YFP; red: blood vessels labelled by SR101). This dataset was warped into a common framework as **(g-i)**. **(g)** Same region as in **(f)**, as obtained by lab X-ray CT. **(h)** Same region as in **(f)**, as obtained by synchrotron X-ray CT. **(i)** Same region as in **(f)**, as obtained by volume EM. **(j-l)** Reconstruction of the complete datasets detailed in **(g-i)**. **(m)** Sketch depicting the extent of the anatomical dataset (blue box), the volume EM contour (pale grey), all functionally imaged ROIs, coloured based on whether they were found inside a glomerulus in EM (green), outside the EM-imaged volume (blue) or located outside of any glomerulus despite being inside the EM-imaged volume (yellow). Detailed reconstructions of the matched EM glomeruli are shown in grey. Centroids of all other glomeruli in the EM dataset that are also in the 2p-imaged volume are shown as filled dots. Centroids of glomeruli in the EM dataset located outside the 2p-imaged volume are shown as empty dots. **(n)** Yield of the correlative experiment, based on number of glomeruli matched across imaging modalities.

## Discussion

Here we have shown that synchrotron X-ray tomography (SXRT) can be used to efficiently link *in vivo* imaging to targeted volume electron microscopy by bridging the gap in accessible length scales. Using SXRT, which provides superior spatial resolution compared to laboratory-based X-ray CT and thus detailed and precise anatomical landmarks, made this pipeline efficient and reliable. We suggest to term this use of X-ray microscopy as “bridging use”^22^. Moreover, we have demonstrated that SXRT provides sufficient resolution to delineate individual cell bodies and at least some neurites in a dense fashion. This in turn provided subcellular context over several millimetres to specific, targeted high resolution volume EM measurements as we have shown for the hippocampus and olfactory bulb tissue. Thus, SXRT can not only provide bridging information but also be employed for “context use”^22^. We have exploited this to show that the trunk dendrites of more superficial CA1 pyramidal neurons display more spines containing spine apparatus in the stratum radiatum than neighbouring dendrites of deeper cells. Deep and superficial CA1 pyramidal neurons display specific spine patterns in their dendritic tufts^44^. Because our sampled region belonged to CA1a, more superficial neurons would receive stronger input from the lateral entorhinal cortex into their tufts and display a larger spine density in their tuft dendrites^44^. Here we have shown that their trunk dendrites also contain more spines with spine apparatus, a specialised form of endoplasmic reticulum involved in the regulation of the dynamics of calcium transients^1,45–48^. Finally, we have shown that combining functional imaging with SXRT and EM **(Fig. 7)** might not only attribute anatomical context for functional imaging but also help to disambiguate functional imaging data itself.

To allow combining *in vivo* imaging with lab and synchrotron-based μCT and ultimately EM, it was critical to obtain sufficient contrast with all modalities as well as to avoid radiation damage. For higher-resolution imaging with SXRT techniques, the radiation dose will have to be considerably increased and cryogenic cooling during data collection might become necessary to mitigate potential radiation damage^33^. A key challenge for volume EM has been to reach sufficient contrast for low current, high-speed imaging, which has been overcome by en-bloc staining protocols rich in heavy metal content. We found that standard osmium/uranium en-bloc EM staining^37^ is not only compatible with SXRT but moreover provides excellent contrast. It offers, however, a limited stain penetration depth, constraining sample sizes in our case to ~600 μm with the reliability of tissue penetration further reduced by prior *in vivo* imaging (not shown). Different variants of reduced osmium/uranium staining protocols^37,54,55^ have been suggested, and future improvements and tailoring of protocols might further optimise the tissue preparation for SXRT imaging, allowing full exploitation of SXRT’s capability to image large samples.

Heavy metal stains as used here limit the size of the samples for SXRT due to their high X-ray absorption cross-section. There are several ways in which this challenge could be overcome. Firstly, the high metal load requirement imposed by serial block-face EM can be mitigated if the charge build-up that takes place during imaging is compensated by other factors^56^, such as conductive resin^57^, focal charge compensation during imaging^58^, or the use of other volume EM imaging techniques that do not impose this restriction (ATUM-mSEM, FIB/SEM). These approaches would all enable volume EM imaging of stained samples with lower (or different) metal content, thereby relaxing the requirements for heavy metal staining. Lighter metal staining combined with X-ray phase-contrast imaging would enable imaging larger tissue blocks with SXRT at similar energy. Furthermore, new and upcoming fourth-generation synchrotron sources^59–62^ offer substantially increased coherent photon flux at high energies facilitating phase-contrast imaging of larger specimens which are highly absorbing at lower energies. While traditional lab-based μCT systems are not capable of producing sufficient density resolution to reliably visualise somata or dendrites, synchrotron X-ray imaging has become more accessible to a wider community in recent years. Tomography beamlines like the ones used here are available at numerous synchrotrons around the world^63^. Moreover, the increased coherent flux of fourth-generation light sources^62^ together with technical advances in e.g. undulator technology will likely make tomography at synchrotrons a rapid, high-throughput technique. Recently developed high-performance laboratory X-ray sources in turn have allowed phase-contrast imaging outside of synchrotrons^24,64^ and could provide another alternative for rapid large volume X-ray imaging at micrometre resolution for broad use.

Here, we have employed synchrotron X-ray CT to image mm^3^ volumes of stained brain tissue with a voxel size of up to 325 nm. These datasets allowed delineating neurites in large tissue samples and provided context for subsequent targeted volume electron microscopy that could reliably identify even smaller structures such as fine dendrites, axons or synapses. Recent developments in coherent X-ray imaging techniques such as near- and far-field ptychography^65,66^ or nano-holotomography^33^ can provide 3D structural information with a spatial resolution approaching tens of nanometres. In principle, this would suffice to reliably detect all structures needed for connectomics analysis^2^. So far, such resolution has not been achieved for soft tissue. However, recent developments in nano-holotomography have demonstrated <100 nm resolution for stained mouse cortical brain samples, demonstrating the potential of coherent X-ray tomography for neuroscience^33^.

Thus, with improvements to data acquisition and analysis, and several synchrotrons recently or about to be upgraded to 4^th^-generation sources^59–62^, synchrotron X-ray imaging is expected to become a tool for rapid large-volume, subcellular anatomical analysis and will help to bridge the current gap in length scales between *in vivo* physiology and high-resolution EM in a routine manner.

## Methods

### Animals

Animals in this study were all 9-12 weeks-old mice. For experiment C525, we used a male transgenic mouse resulting of MOR174/9-eGFP^67^ crossed into M72-IRES-ChR2-YFP^68^ (JAX stock #021206) crossed into a Tbet-cre driver line^69^ (JAX stock #024507) crossed with a GCaMP6f reporter line^70^ (JAX stock #028865). For experiment C432, we used a female transgenic mouse carrying the latter three constructs. For the rest of experiments, we used male mice of C57Bl/6 background. All animal protocols were approved by the Ethics Committee of the board of the Francis Crick Institute and the United Kingdom Home Office under the Animals (Scientific Procedures) Act 1986.

### *In vivo* imaging

#### Surgical and experimental procedures

Prior to surgery all utilised surfaces and apparatus were sterilised with 1% trigene. Mice aged 10 weeks were anaesthetised using a mixture of fentanyl/midazolam/medetomidine (0.05 mg/kg, 5 mg/kg, 0.5 mg/kg respectively). Depth of anaesthesia was monitored throughout the procedure by testing the toe-pinch reflex. The fur over the skull and at the base of the neck was shaved away and the skin cleaned with 1% chlorhexidine scrub. Mice were then placed on a thermoregulator (DC Temperature Controller, FHC, ME USA) heat pad controlled by a temperature probe inserted rectally. While on the heat pad, the head of the animal was held in place with a set of ear bars. The scalp was incised and pulled away from the skull with four arterial clamps at each corner of the incision. A custom head-fixation implant was attached to the base of the skull with medical super glue (Vetbond, 3M, Maplewood MN, USA) such that its most anterior point rested approximately 0.5 mm posterior to the bregma line. Dental cement (Paladur, Heraeus Kulzer GmbH, Hanau, Germany; Simplex Rapid Liquid, Associated Dental Products Ltd., Swindon, UK) was then applied around the edges of the implant to ensure firm adhesion to the skull. A craniotomy over the left olfactory bulb (approximately 2 x 2 mm) was made with a dental drill (Success 40, Osada, Tokyo, Japan) and then immersed in ACSF (NaCl (125 mM), KCl (5 mM), HEPES (10 mM), pH adjusted to 7.4 with NaOH, MgSO4.7H2O (2 mM), CaCl2.2H2O (2 mM), glucose (10 mM)) before removing the skull with forceps. The dura was then peeled back using fine forceps. A layer of 2% low-melt agarose diluted in ACSF was applied over the exposed brain surface before placing a glass window cut from a cover slip (borosilicate glass #1 thickness [150 μm]) using a diamond scalpel (Sigma-Aldrich) over the craniotomy. The edges of the window were then glued with medical super glue (Vetbond, 3M, Maplewood MN, USA) to the skull.

Following surgery, mice were placed in a custom head-fixation apparatus and transferred to a two-photon microscope rig along with the heat pad. The microscope (Scientifica Multiphoton VivoScope) was coupled with a MaiTai DeepSee laser (Spectra Physics, Santa Clara, CA) tuned to 940 nm (<30 mW average power on the sample) for imaging. Images (512 x 512 pixels, field of view 550 μm x 550 μm) were acquired with a resonant scanner at a frame rate of 30 Hz using a 16x 0.8 NA water-immersion objective (Nikon). Using a piezo motor (PI Instruments, UK) connected to the objective, a volume of ~300 μm was divided into 12 planes resulting in an effective volume repetition rate of ~2.5 Hz. The odour port was adjusted to approximately 1 cm away from the ipsilateral nostril to the imaging window, and a flow sensor (A3100, Honeywell, NC, USA) was placed to the contralateral nostril for continuous respiration recording and digitized with a Power 1401 ADC board (CED, Cambridge, UK).

Following odour stimulations (see below), Sulforhodamine 101 (Sigma Aldrich, 100 μM final concentration) was injected intraperitoneally to label blood vessels and a z stack of the entire volume was acquired, consisting of 88 images each covering (464 μm)^2^ with (358 nm)^2^ pixels, separated in z by 5 μm **(Table 1)**.

**Table 1.**
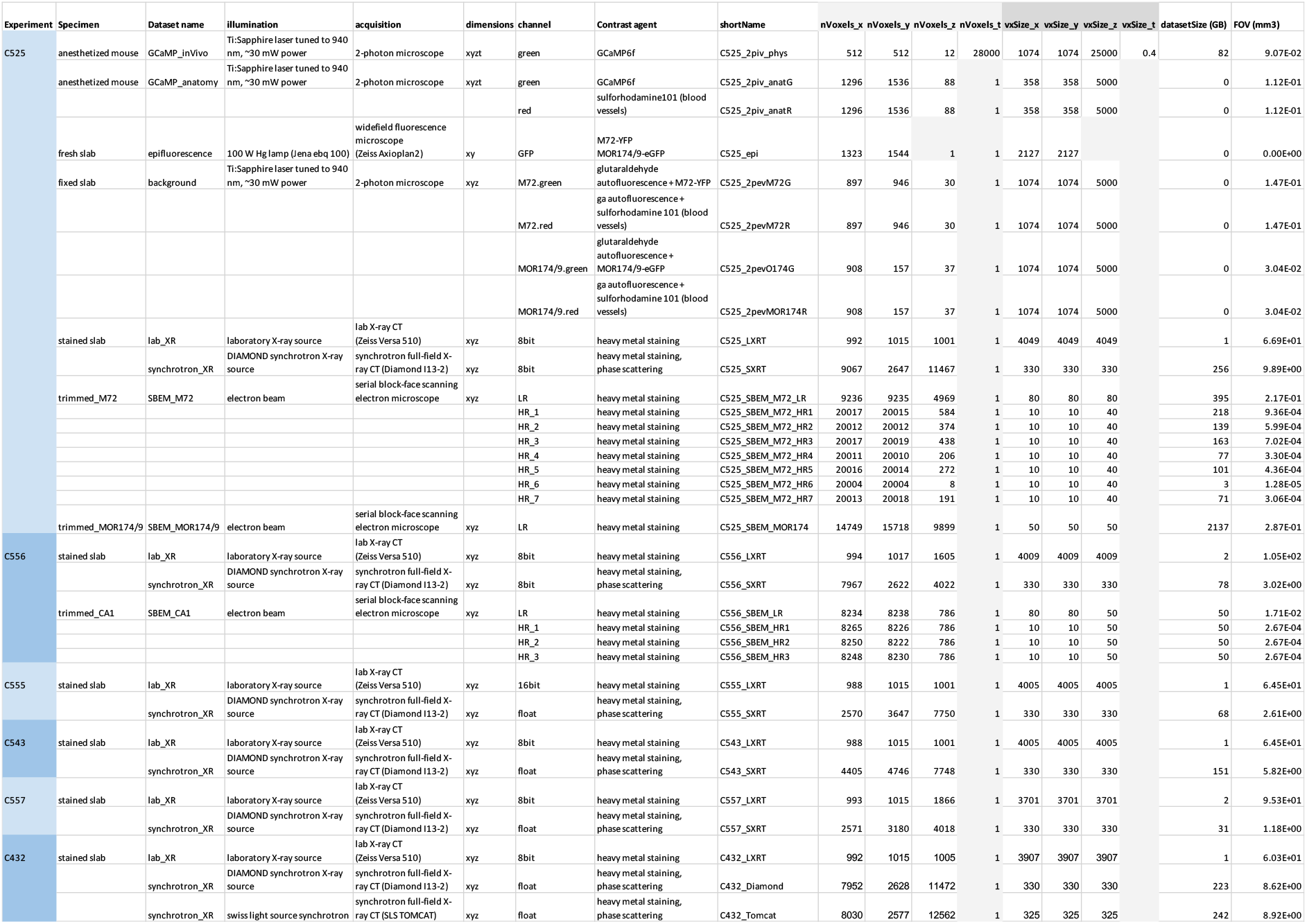
Datasets. Details of all datasets of correlative multimodal imaging experiments described in this study.

#### Odour stimulation

Odour stimuli were delivered using a set of custom-made 6-channel airflow dilution olfactometers. In brief, volumes of 3 ml of a set of 48 monomolecular odorants (Sigma-Aldrich, St. Louis MO, USA) were pipetted freshly for each experimental day into 15 ml glass vials (27160-U, Sigma-Aldrich, St. Louis MO, USA). Odours were diluted 1:20 in air before being presented to the animal at 0.3 litres/min using custom Python Software (PulseBoy; github.com/RoboDoig). Odours were prepared for 3 seconds in tubing before a final odour valve was triggered to open at the beginning of an inhalation cycle. During anaesthetised recordings all stimuli were presented with a 30 s inter-stimulus interval. To minimise contamination between odour presentations, a high flow clean air stream was passed through the olfactometer manifolds during this time.

#### Data analysis

Motion correction, segmentation and trace extraction were performed using the Suite2p package (https://github.com/MouseLand/suite2p). Regions of Interest (ROIs) corresponding to glomeruli were manually delineated based on the mean fluorescence image in Fiji^71^. The fluorescence signal from all pixels within each ROI was averaged and extracted as a time series with ΔF/F = (F-F0)/F0, where F = raw fluorescence and F0 = median of the fluorescence signal distribution. Further analysis was performed with custom written scripts in Matlab.

### Tissue preparation

#### Dissection

Mice were sacrificed and 600 μm-thick slabs of brain areas of interest were sliced in ice-cold dissecting buffer (phosphate buffer 65 mM, 0.6 mM CaCl2, 150 mM sucrose) with an osmolarity of 300 ±10 mOsm/L using a LeicaVT1200 vibratome and immediately transferred to ice-cold fixative (2% glutaraldehyde in 150 mM sodium cacodylate buffer pH 7.40, 300 mOsm/L). A stereoscope (Leica MZ10 F) was mounted above the vibratome to guide slicing and confirm presence of fluorescent landmarks on the fresh slabs when present. At this point, the olfactory bulb slab was quickly imaged using a widefield fluorescence microscope (Zeiss Axioplan2), recording the coarse location of the fluorescently-tagged glomeruli M72-YFP and MOR174/9-eGFP. Samples were left in the same fixative overnight, at 4°C. The fixative was then washed with wash buffer (150 mM sodium cacodylate pH 7.40, 300 mOsm/L) three times for 10 minutes at 4°C. The OB slab was then imaged at the 2-photon microscope to record the 3D anatomical background autofluorescence generated by glutaraldehyde, in the locations surrounding both tagged glomeruli. Overall, samples were kept in an ice-cold, osmolarity-checked buffer.

#### Staining, dehydration and embedding

Slabs were stained with heavy metals using an established ROTO protocol^37^ using an automated tissue processor (Leica EMTP). Briefly, they were first stained with reduced osmium (2% OsO4, 3% potassium ferrocyanide, 2mM CaCl2 in wash buffer) for 2h at 20°C, followed with 1% thiocarbohydrazide (aq) at 60°C for 50 min, 2% osmium (aq) for 2h at 20°C, and 1% uranyl acetate (aq) overnight at 4°C. On the next day, the samples were further stained with lead aspartate (prepared as in ^72^) for 2h at 60°C. Samples were washed with double-distilled water six times for 10 minutes at 20°C between each staining step, except warmer washes before and after TCH (50°C) and before LA (60°C).

Samples were then dehydrated with increasing ethanol solutions (75%, 90%, 2×100%), transferred to propylene oxide, and infiltrated with hard epon^73^ mixed with propylene oxide in increasing concentrations (25%, 50%, 75%, 2×100%). Finally, samples were polymerised individually into plastic moulds for 72h at 70°C.

### SXRT imaging

Samples were imaged with full-field X-ray tomography at the I13-2 beamline of the Diamond synchrotron (Didcot, UK) using a polychromatic X-ray beam from an undulator source, filtered by 1.34 mm pyrolytic graphite, 2.1 mm aluminium, 0.035 mm silver, and 0.042 mm palladium. This produced an X-ray spectrum with an effective energy of approximately 22 keV. Samples were mounted at a distance of 52 mm upstream of the detector system. The latter consisted of a scintillation screen (34 mm-thick Europium-doped Gadolinium Gallium Garnet (GGG:Eu)), a 10x objective lens, 2x tube lens and a pco.edge 5.5 camera (sCMOS chip with 2560 x 2160 pixels). This system yielded an effective pixel size of 330 nm.

A set of 2-24 local tomography scans (with an overlap of 0.22 mm in the horizontal and 0.1 mm in the vertical direction) was acquired for each specimen to cover a total volume of 1.2-9.9 mm^3^**(Table 1)**. For each tomography, 3001 projections were recorded at equi-distant viewing angles over 180° of sample rotation with an exposure time of 0.4 s per frame, which resulted in a total scan time of approximately 20 minutes per tomography dataset. Phase tomograms were reconstructed by applying a commonly used single-distance phase retrieval algorithm^74^ to the 2D projections and subsequently performing the tomographic reconstruction using a filtered back-projection algorithm. For the single-distance phase retrieval, a ratio of the real part delta and imaginary part beta of the refractive index was set to delta/beta = 1, based on visual optimisation of the reconstructed images.

One sample (C432) was subsequently imaged with full-field X-ray tomography at the TOMCAT beamline of the Swiss Light Source at the Paul Scherrer Institut (Villigen, Switzerland) using a multilayer monochromatic X-ray beam filtered by 0.1 mm aluminium and 0.01 mm iron. This produced an X-ray spectrum with an effective energy of 21 keV. Samples were mounted at a distance of 50 mm upstream of the detector, consisting of a scintillation screen (LuAg:Ce 20um), a 20x objective lens and a pco.edge 5.5 camera, yielding to an effective pixel size of 325 nm. 24 tomography scans (with an overlap of 0.25 mm in the horizontal and 0.03 mm in the vertical direction) were acquired covering the entire specimen (8.9 mm^3^). For each tomography, 2601 projections were recorded over 180° of sample rotation with an exposure time of 0.3 s per frame, which resulted in a total scan time of approximately 13 minutes per tomogram. The tomograms were reconstructed using the same protocol as above. The parameters used for the single-distance phase retrieval were put to delta = 2e-6 and beta = 6e-7. An unsharp mask filter was applied with a stabilizer of 0.6 and a gaussian kernel filter of 1.0. The tomographic reconstruction was performed using the GridRec algorithms^75^.

The different volumes obtained by the local tomographies were stitched together using a 3D non-rigid stitching algorithm based on local pairwise cross-correlation^76^.

### EM imaging

Three serial block-face electron microscopy (SBEM) datasets were obtained for this study, to explore in detail the olfactory bulb (C525a,b) and the hippocampus (C556) of samples previously imaged with other imaging modalities.

In all cases, the parent stained sample was scanned with lab X-ray CT (Zeiss Versa 510) and a trimming plan was designed to render the regions of interest optimally accessible for the SBEM. Additional datasets obtained from other modalities (2p *ex vivo* datasets of regions containing genetically-tagged fluorescent glomeruli) were also warped to the same scene in some cases (C525a,b and C432) to determine the ROI **(Supp. Fig. 6a,b)**.

Several lab X-ray CT datasets taken at different steps of trimming were warped onto the same scene iteratively (using BigWarp^77^) **(Supp. Fig6 c1-c2, d1-d2)**, guiding the trimming process and ensuring the ROI was preserved at all times. The trimming process involved the use of tools of increasing precision as required: hand-held single-edge blades, ultramicrotome-guided glass knives and a 4mm diamond knife (Leica UC7 ultramicrotome, Diatome histo knife). The final sample was a <1mm^3^ tissue cuboid with a diamond knife-polished surface displaying some tissue. The sample sides were then covered with a silver colloidal suspension (SPI supplies), and the sample was sputter-coated with 10 nm platinum (Quorum Q150R S).

All serial block-face EM datasets were obtained with single images covering the desired field of view. Datasets C525a and C556 were acquired using a 3View2 – variable pressure Zeiss Sigma serial block-face SEM using focal charge compensation^58^. In both cases, datasets were acquired using a 2.5 kV electron beam and a 30 μm aperture, and multiple ROIs were acquired at high and low resolution covering narrower and wider fields of view, respectively **(Table 1)**. In particular, C525a contained 6004 slices across 11 concatenated datasets in z. Seven high-resolution regions (acquired in z-order 1,2,3,5,7,9,11) were composed altogether by 2078 images acquired at 2.0 μs/pixel, each covering (200 μm)^2^ with (10 nm)^2^ pixels, separated in z by a slice thickness of 40 nm. A reconstructed continuous low-resolution dataset contained 4970 images acquired at 0.5 μs/pixel covering (737 μm)^2^ with (80 nm)^2^ pixels, separated in z by a slice thickness of 80 nm. Acquiring the C525a datasets took 25 days. C556 was composed by one low-resolution dataset and three high-resolution datasets from narrower regions in the same volume, all containing 787 images separated in z by a slice thickness of 50 nm. The images of the low-resolution dataset were acquired at 0.5 μs/pixel and covered (655 μm)^2^ with (80 nm)^2^ pixels. The images of the high-resolution dataset were acquired at 2.0 μs/pixel and covered (82 μm)^2^ with (10 nm)^2^ pixels. Acquiring the C556 datasets took 5 days. C525b was acquired using a 3View2 – Zeiss Merlin serial block-face SEM under high vacuum, a 2.5 kV electron beam with a landing current of 0.5 nA and an OnPoint backscattered electron detector. The dataset was composed by 9899 images acquired at 0.5 μs/pixel covering (700 μm)^2^ with (50 nm)^2^ pixels, resulting in a dose of 0,62e^-^/nm^2^, separated in z by a slice thickness of 50 nm. Acquiring the C525b dataset took 18 days.

Additionally, the micrographs shown in Supp. Fig. 3 a-b were taken from C046, a serial blockface EM dataset previously reported in reference ^78^.

#### Image registration

Each serial block-face EM dataset was registered using Voxelytics Align (scalable minds, Potsdam, Germany), which uses pairwise feature matching between adjacent slices, RANSAC inlier detection and global relaxation to create mesh-tessellated affine transformations. When a number of slices had to be discarded, the closest non-discarded slices were duplicated in the final dataset thereby keeping the total number of slices unaltered.

C525a datasets **(Supp. Fig5 a2)** were registered by removing 3 slices **(Supp. Fig5 a4)**. C556 datasets were registered without any slice loss **(Supp. Fig5 a5)**.

For C525b **(Supp. Fig5 a1)**, 595 slices (6%) had to be discarded to register the dataset **(Supp. Fig5 a3)**, generating one single continuous gap of 10 slices (0.5 μm) and an average gap size of 2.7 slices (135 nm). These slices were detected by looking at maximum variation and at minimum matched features between any two adjacent slices. These artefact detections and the registration were run on a masked version of the dataset only containing tissue voxels. This mask was essential to avoid picking up statistics from imaging artefacts present in areas filled with resin (such as charging artefacts). The final step involved correcting for lateral drift across z, which is most likely a registration artefact arising from the presence of features in the dataset that have a preferred orientation.

### Data processing of SXRT and EM data

Datasets were stored as a chunked file format with a resolution pyramid (wkw) and explored in webKnossos^39^.

#### Warping

Linked datasets obtained with different imaging modalities were warped into a common framework using BigWarp^77^**(Supp. Fig. 5b)**. Conserved landmarks were manually identified and a first affine transformation of the moving dataset was generated allowing to find further local paired landmarks between both datasets. This iterative process of landmark deposition and dataset transformation was performed until no observable improvement in overlap was found, using 11-171 paired landmarks. The number of landmarks depended on the modalities being warped: 27-53 landmarks were used for warping 2-photon datasets of fixed tissue into a lab X-ray CT **(Supp. Fig. 5 b1-b3, b8)**; 11-25 to warp lab and synchrotron X-ray CT **(b4-b5, b8)**, 50-51 to warp synchrotron X-ray CT and volume EM, and 171 to warp *in vivo* 2-photon to volume EM **(b6-b7, b9)**. The nature of landmarks also depended on the modalities being warped, since this relies on conserved features across both modalities: glomerular contours and tissue boundaries were features present in all datasets, blood vessels were well defined in the *in vivo* 2-photon dataset, and these and nuclei were easily found in synchrotron X-ray CT and volume EM. This set of landmarks was then used to generate a warp field to move between both datasets. The Bigwarp engine was integrated into Matlab using custom code and integrated with a webKnossos skeleton analysis toolbox^79^. We constructed a graph with nodes representing datasets and edges representing transform functions, either involving spline transformations as introduced above or scale-offset transformations. This graph contained all information required to transfer annotations from the coordinate system of a dataset to the coordinate system of an arbitrary linked dataset. This allowed warping skeletons between any connected pair of datasets, a key tool for the comparative analyses presented. In some cases, the image data from regions of interest spanning part or the whole moving dataset were warped into the target dataset’s space and rendered in a common viewport using Amira (Thermo Fisher) for illustrative purposes.

### Data analysis of SXRT and EM data

#### Apical dendrite tracing and SXRT traceable length analyses

Mitral cell apical dendrites were traced in EM and SXRT datasets. Somata of mitral cells were identified based on their histological and morphological features^80^, mainly a large cell body located in a monolayer parallel to the glomerular layer, 200-400 μm below it, with a large, pale nucleus containing a prominent nucleolus and surrounded by abundant cytoplasm. All mitral cell somata (372) in the EM dataset (C525b) were manually tagged by creating a node in their nucleus and their names and coordinates were stored in a webKnossos skeleton file (*.nml). 50 somata were randomly selected without replacement for this analysis and saved as a new skeleton. This file was warped onto the SXRT dataset space using the integrated warping toolbox. This enabled manually tracing the same cell in the native version of each imaging modality. Three trained tracers were assigned the task of manually tracing all apical dendrites in both EM and SXRT datasets. Briefly, it involved identifying the apical dendrite that arises from the cell body of the seeded nucleus, and then tracing that apical dendrite centrifugally until reaching the glomerulus without creating any branches. Once inside the glomerulus, the most prominent dendrite was to be traced as well. All EM skeletons (with 3 traces per neuron) were pooled and manually curated (all could be reconstructed: 34/50 were correctly traced by all, 16/50 were correctly traced by two tracers). The longest tree of the correctly traced ones was proofread and chosen as consensus apical dendrite for that one neuron, generating ground truth apical dendrite skeletons for all 50 mitral cells. These ground truth skeletons were warped into the SXRT space using the integrated warping toolbox described above. For each tracer, the apical dendrites traced in the SXRT dataset were paired with the warped ground truth (EM) apical dendrite of the same cell. Both trees would start at the same position (nucleus).

A polynomial cubic spline function (Matlab File Exchange) was fitted to both trees, and evenly spaced nodes were created every 320 nm. For every node in the SXRT-traced tree, the Euclidean distance to the closest node in the warped EM tree was calculated and plotted as a function of the position in the SXRT-traced tree. These distances displayed a bimodal distribution with the two peaks arising from correctly and incorrectly traced dendrites respectively. A tree was considered “lost” when the closest distance to the consensus tree was >12μm, and the distance from the somatic seed to the point at which it was lost was defined as the correctly traced distance. Most tree losses took place within the first 40 μm of dendrite, which can be attributed to the beginning of the apical dendrite. We named this challenge as the “linkage” between the nucleus node and the apical dendrite. We called “linked” all trees that were correctly traced for at least 35 μm. This enabled splitting all trees between kept or lost, linked or not linked, or shorter than linked and kept thresholds.

A similar approach was used in the hippocampus. In this experiment, the EM dataset was configured so the xy plane matched the direction of CA1 apical dendrites and the z axis evolved almost perpendicular to the pyramidal layer. A bounding box of 180*200 μm^2^ in x,y and spanning the entire z of the dataset (39 μm) was defined so it would contain the central CA1 region in the dataset. Pyramidal neurons were easily identifiable based on their large soma with a nucleolus and condensed chromatin inclusions located in a packed layer with a clearly distinguishable apical dendrite arising from the cell body. All 175 CA1 pyramidal somata in the bounding box were tagged by creating a node in their nucleus. One trained tracer was assigned to trace all their dendrites in both EM and SXRT datasets. In EM, the dendrites of 132 somata could be traced, generating ground truth apical dendrite skeletons for these 132 CA1 pyramidal neurons. 90 of these could be traced in the SXRT data. In this case, a warped version of the SXRT dataset on the EM space was also used to complement the tracings when helpful. The distances between SXRT and EM trees also displayed a bimodal distribution, which was used to set the “lost” threshold to 6 μm. The “linkage” threshold was set to 35 μm.

The same tracing and analysis methods were also used to compare apical dendrites of OB mitral cells traced in the two different microtomography beamlines **(supp. Fig. 4)** (as in previous OB analyses: interpolation step = 320nm, lost threshold = 12 μm). Here, I13-2 tracings were used as references and TOMCAT tracings were evaluated against them. Since the distance analysis only applies to dendrite tracing pairs, only cells traced in both datasets were analysed (33 out of 50 randomly seeded mitral cell somata, the remaining 17 were traceable in only one or neither dataset).

#### Correlative SXRT-EM analyses of hippocampal CA1 neurons

##### Soma depth of CA1 pyramidal neurons

A linear regression line was fitted to the x,y coordinates of all seeded CA1 pyramidal neuron somata. Soma depth was quantified by the Euclidean distance between a soma and the regression line. Here, the soma z coordinate was ignored as apical dendrites were approximately parallel to the xy image plane. Deeper cells have more positive scores. A subset of the 90 CA1 apical dendrites traced in the SXRT dataset passed through the field of view of the high-resolution EM dataset. Among them, seven dendrites with varying soma depths were chosen for further spine and spine apparatus analyses.

##### Dendritic spine tracing and spine apparatus tagging

By warping SXRT apical dendrite tracings into the high-resolution EM dataset space, dendrites corresponding to the seven target cells were identified. Similar to apical dendrite tracings, one trained tracer manually annotated all dendrite branches, dendritic spines and spine apparatuses on webKnossos. For each of the seven dendrites, a linear tracing delineating the dendritic shaft was created first. Spine-level annotations were added in a second step. When a spine was encountered, a node was placed within the spine head which ultimately connected to a node inside the spine neck. To ensure accurate spine distribution analyses, this spine branch was extended to intersect perpendicularly with the dendritic shaft skeleton and was merged into it. For spines with more than one spine apparatus cisterns inside its head or neck, the spine head node was commented to indicate the presence of spine apparatus.

##### Density and distribution analyses of spines and spine apparatus

Once the spine-level dendrite tracing was finalised, the traced skeleton was split into individual branchlets and grouped as trunk or apical obliques. For each individual dendrite branchlet, the number of spines and spines with spine apparatus were counted. The dendritic spine count was based on the number of spine necks joining the dendritic shaft, and each spine also carried a binary present/absent spine apparatus score. For example, a branched spine with spine apparatuses in both of its heads was counted as one spine with spine apparatus. Both interspine distance and inter-spine apparatus distance were measured as distances between intersecting points of adjacent spines on the dendritic shaft. In total from the seven cells, 75 dendrite branchlets, which were 2 mm in length in total, were inspected and their 3019 spines and 417 spine apparatuses analysed for density and distribution.

Spine density was defined as the total number of spines divided by the total shaft length pooled across all dendrite branchlets of the same order for each cell.

Spine apparatus density was similarly defined as the total number of spines possessing a spine apparatus divided by the total length of dendritic shaft pooled.

If spines emerged from the shaft randomly, the cumulative distribution of inter-spine distances should follow a Poisson distribution *f*(*x*) = *λe*^-*λx*^, where *λ* equals the average inter-spine distance. For each cell, this null hypothesis was tested using one-sample Kolmogorov-Smirnov test. p-values < 0.05 were considered significant. Similarly, spine apparatus may also occur randomly in spines. Therefore, a two-sample Kolmogorov-Smirnov test was used to compare the inter-spine distance distribution against the inter-spine apparatus distribution for apical oblique dendrite branchlets of each cell. For visualisation, cumulative distribution of ISD and ISAD were normalised by their mean values and plotted together with the cumulative distribution of a unit exponential function (*λ* =1). Lastly, all density and distribution characteristics were analysed in light of soma depths by plotting the scores against soma depth.

##### Most distant nearest spines

When comparing the cumulative distribution of ISD to the null hypothesis (i.e. Poisson distribution), we observed that the ISD rise above the null hypothesis at large ISD distances consistently in all seven cells for both trunk and apical oblique dendrites (supp. Fig. 8a1 & a2). One potential explanation is that there is a minimum spine density that caps the ISD value. To test this hypothesis, we found the maximum ISD value for each dendrite branchlet, focusing on apical oblique dendrites only because those traces contained more spines. Since observed ISD cannot be longer than the length of a sample dendrite, any bias introduced by dendrite length needs to be eliminated. Therefore, maximum ISD values were plotted against the length of each sampled dendrite branchlet (supp. Fig. 8c). In total, 53 out of 58 apical oblique dendrites contained more than one spine (i.e. at least one ISD value) and were included in the analysis.

## Acknowledgements

We are grateful to the advanced light microscopy, biological research, electron microscopy and scientific computing science technology platforms of the Francis Crick Institute and thank Kevin Briggman and Sina Tootoonian for discussion.

This research was funded in whole, or in part, by the Wellcome Trust (FC001153). For the purpose of Open Access, the author has applied a CC BY public copyright licence to any Author Accepted Manuscript version arising from this submission. This work was carried out with the support of Diamond Light Source, beamline I13-2 (proposal 20274) and the TOMCAT beamline of the Swiss Light Source at the Paul Scherrer Institut (proposal 20190417). This work was supported by the Francis Crick Institute, which receives its core funding from Cancer Research UK (FC001153), the UK Medical Research Council (FC001153), and the Wellcome Trust (FC001153); by the UK Medical Research Council (grant reference MC_UP_1202/5), and a DFG postdoctoral fellowship to TA. ATS and TWM are Wellcome Trust Investigators (110174/Z/15/Z, 214333/Z/18/Z). AP has received funding from the European Research Council under the European Union’s Horizon 2020 Research and Innovation Programme (grant no. 852455).

## Supplementary Figures

**Supp. Fig. 1.**
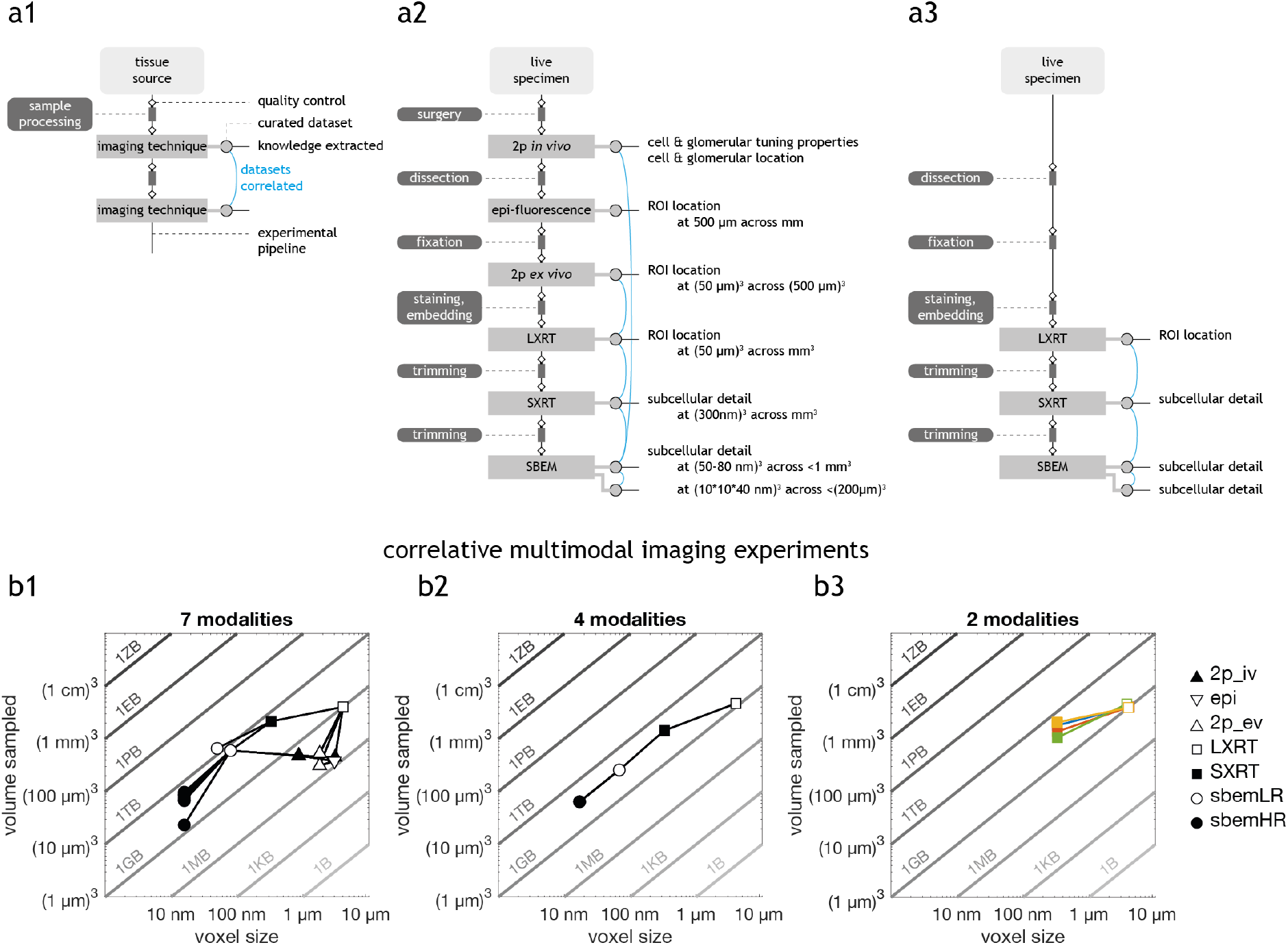
Comparison of correlative multimodal imaging (CMI) pipelines. **(a)** Flowchart diagram **(a1)**, applied to a CMI pipeline including 6 imaging techniques **(a2)** and to a CMI pipeline including 3 imaging techniques **(a3)**. This diagram shows the elements that might affect the throughput of a CMI pipeline and is helpful for identifying bottlenecks. Samples are processed before every imaging technique to match compatibility requirements and enhance signal detection. A quality control step is present before any sample preparation and imaging step, aimed to maximise the success rate of the following steps. Raw data has to be processed to generate a curated dataset. Curated datasets can be correlated, thereby enabling to link the information obtained in their respective analyses and augmenting the knowledge extracted from the pipeline. **(b)** Correlated datasets in a CMI experiment including 7 modalities **(b1),** 4 modalities **(b2)**, and in four experiments including the same 2 modalities each **(b3)**. Note that some imaging techniques can image the same specimen at different sampling rates, providing more than one modality (e.g. SBEM high vs low-resolution). Furthermore, some techniques allow imaging multiple regions of interest, thereby providing more than one dataset per modality (e.g. SBEM high-resolution datasets in **b1**). For each dataset, the x axis shows the “voxel size” being the cubic root of the product of the acquired voxel size in x,y,z – thereby representing the length of the side of a voxel if voxels were isotropic. The y axis shows total volume sampled. In this diagram, datasets containing the same number of voxels are distributed along a diagonal. For reference, diagonals hosting datasets of each order of magnitude are shown in shaded greys and their sizes are indicated assuming they are uncompressed 8-bit images. The dataset marker represents the imaging modality: *2p_iv*, 2-photon *in vivo* Ca^2+^; *epi*, epifluorescence of the dissected slab; *2p_ev*, 2-photon *ex vivo* imaging of the fixed slab; *LXRT*, lab-source X-ray CT; *SXRT*, synchrotron-sourced X-ray CT; *sbemLR* and *sbemHR*, serial block-face electron microscopy at low and high resolution respectively. Datasets warped to each other are linked with an edge. All plots represent CMI pipelines reported in this study. **(b1)** and **(b2)** contain one single experiment each. In **(b3)**, datasets belonging to each experiment are shown in the same colour.

**Supp. Fig. 2.**
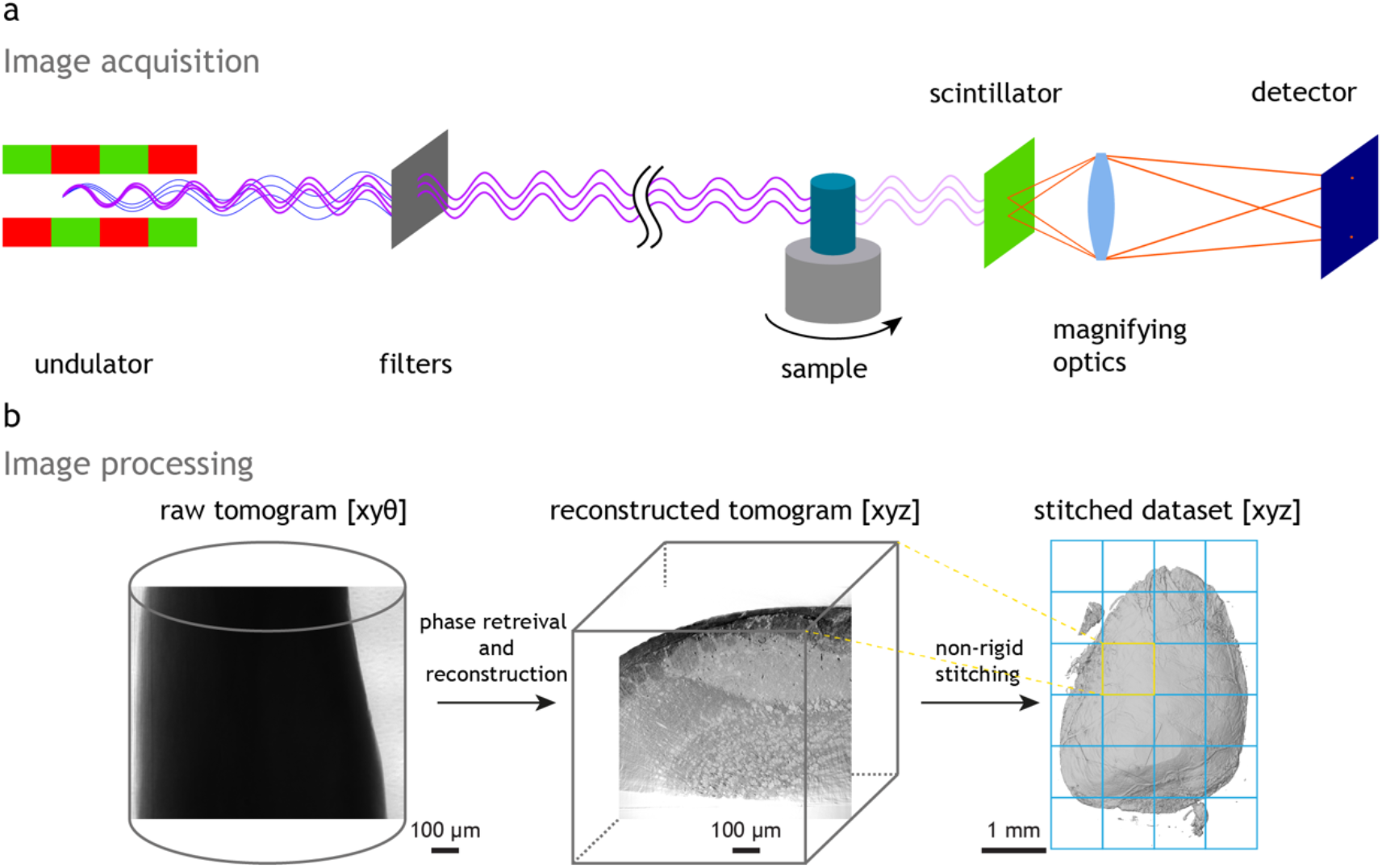
Data acquisition and processing in a synchrotron X-ray CT experiment. **(a)** Simplified schematic of the synchrotron X-ray CT experiment at Diamond I13-2. The undulator (alternating magnetic structures, depicted as red/green) creates polychromatic X-rays from the stored electron beam. X-ray filters remove the lower energy UV and X-ray radiation and only leave the tailored X-ray spectrum for the experiment. The experimental station itself is situated about 230 m downstream of the undulator, and the sample is mounted on a rotation stage. After interacting with the sample via absorption and refraction, the X-rays reach a scintillator crystal which converts them to visible light. Magnifying optics project an image of the scintillator on the chip of the sCMOS camera. The images are then further processed and reconstructed. **(b)** Simplified schematic of the image processing required to generate datasets from the synchrotron X-ray CT experiment. The sample is rotated and a projection of the sample on the detector is recorded at each angular step θ, thereby generating a x,y,θ dataset. This is subsequently reconstructed into a x,y,z dataset. The sample is moved so that overlapping 3D tiles are captured covering the entire region of interest of the sample. The reconstructed x,y,z tiles are stitched together using a non-rigid algorithm to ensure a smooth transition at the boundaries. The stitched dataset represents an x,y,z volume covering the entire region of interest.

**Supp. Fig. 3.**
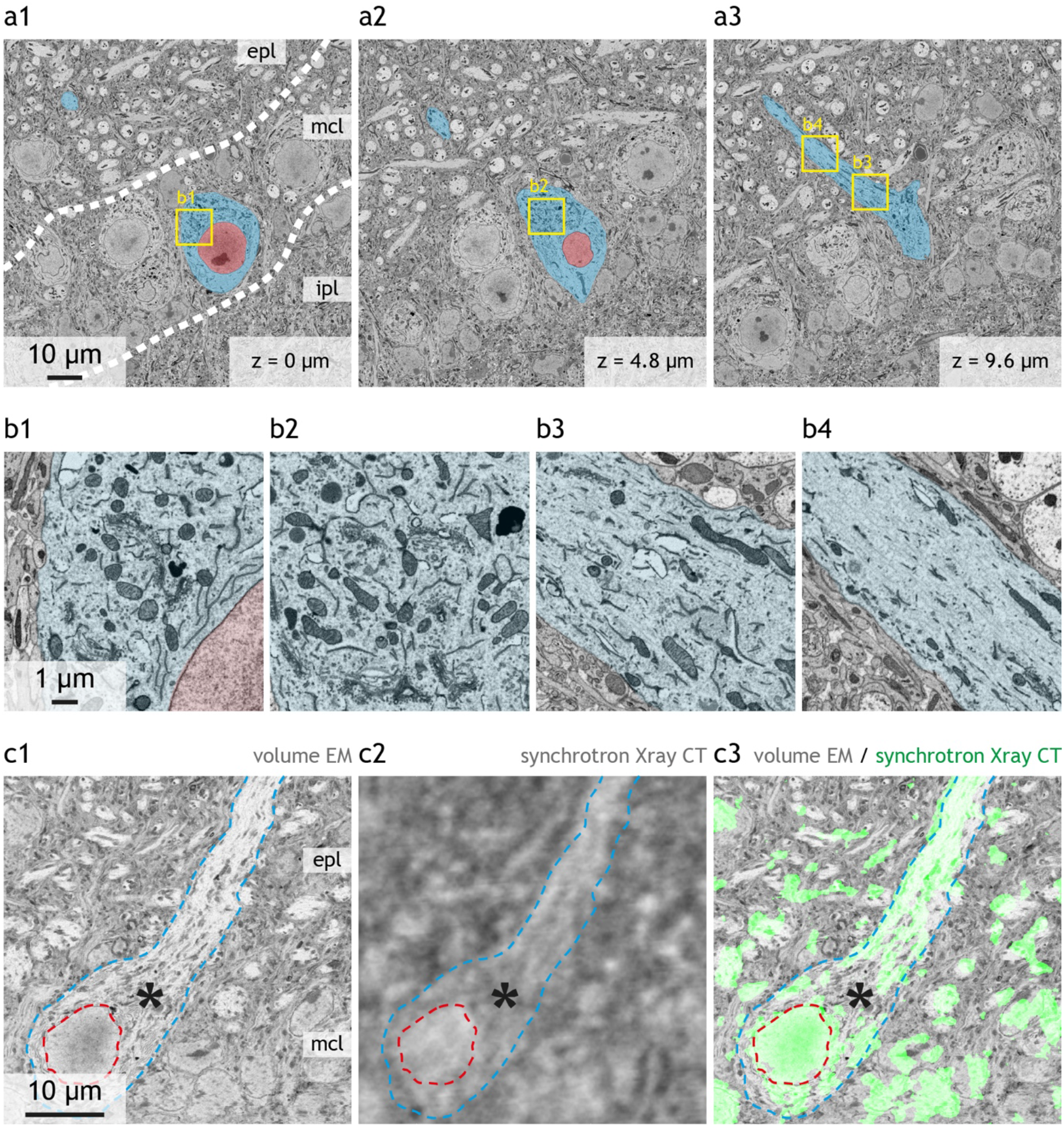
Initial segment of the apical dendrite of a mitral cell. **(a)** Mitral cell (nucleus in red, cytoplasm in blue) shown at three z depths **(a1-3)**. Details of the cytoplasm are shown for four regions: in soma, proximal **(b1)** and distal to nucleus **(b2)**, beginning of apical dendrite **(b3)** and apical dendrite **(b4)**. Note the abundance of electron-dense features in **(b1-3)** – including mitochondria, rough and smooth endoplasmic reticulum, and Golgi apparatus – and how they become much less prominent in **(b4)**. **(c)** Correlative EM **(c1)** and synchrotron X-ray CT **(c2)** of the same region containing a mitral cell. Both datasets were warped to a common space. Nucleus (red dashed line) and cell boundaries (blue dashed line) are labelled, the latter delineating the continuum cell body – apical dendrite. The overlay of both modalities is shown in **(c3)**, with the synchrotron X-ray image thresholded so only data from poorly absorbing regions (e.g. dendrites) is displayed (green channel). Note that in the synchrotron X-ray dataset both the nucleus and the apical dendrite are well defined whereas the initial segment of the apical dendrite can get blended into the surrounding neuropil (asterisk). *epl*, external plexiform layer; *mcl*, mitral cell layer; *ipl*, inner plexiform layer The dataset from which the micrographs in **(a,b)** are taken was previously reported in reference ^78^.

**Supp. Fig. 4.**
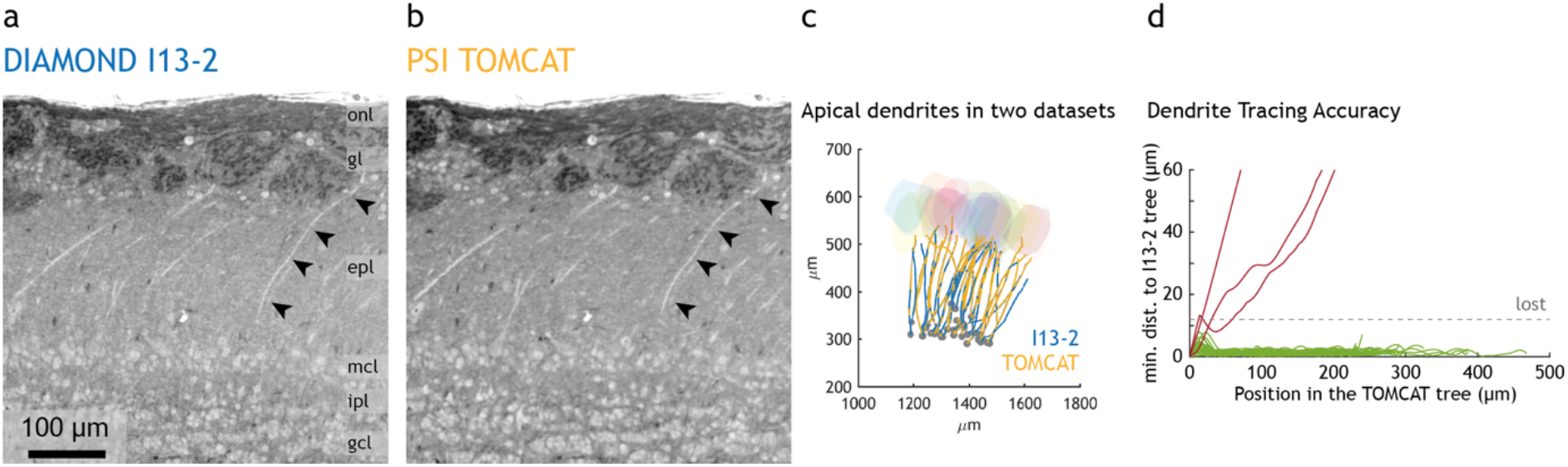
Apical dendrites of mitral cells in synchrotron X-ray CT data acquired at different synchrotron X-ray CT beamlines. **(a-b)** Virtual slice across the same sample region resolved by synchrotron X-ray tomography performed at the Diamond beamline I13-2 **(a)** and at the SLS beamline TOMCAT **(b)**. **(c)** The apical dendrites of the same mitral cells were traced independently in both datasets and subsequently warped and plotted in a common space. Overlap between paired traces indicates a good agreement between the same dendrites traced in the two datasets. The rough contours of glomeruli are shown as coloured blobs. **(d)** Distance between traces of paired dendrites, analysed as in **Fig. 3 b, e.** 91% of the dendrites were accurately traced in both datasets (30 out of 33 dendrites traced in both datasets). *onl*, olfactory nerve layer; *gl*, glomerular layer; *epl*, external plexiform layer; *mcl*, mitral cell layer; *ipl*, inner plexiform layer; *gcl*, granule cell layer.

**Supp. Fig. 5.**
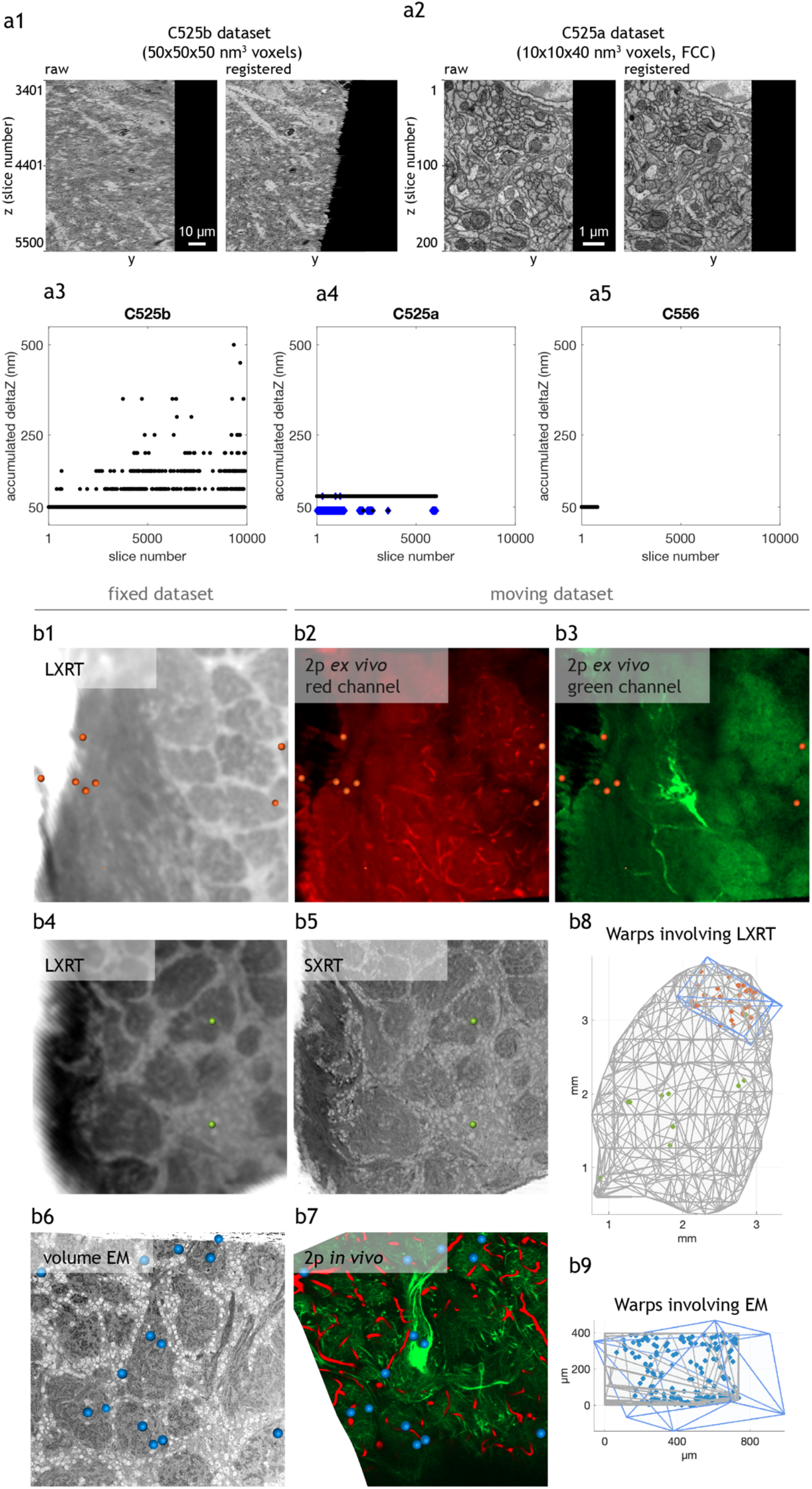
Image registration for serial block-face electron microscopy datasets, and warping datasets from different modalities using manually seeded landmarks. **(a)** Registered outcomes from three experiments involving serial block-face EM. **(a1)** Unregistered (left) and registered (right) yz reslice of the same region in the C525b dataset. **(a2)** Unregistered (left) and registered (right) yz reslice of the same region in the C556 dataset. **(a3-a5)** Summary plots displaying the z distance between every two consecutive slices in the curated dataset, for datasets in C525b **(a3)**, C525a **(a4)** and C556 **(a5)**. In C525a **(a4)**, the experiment consisted of several high-resolution datasets spaced 40 nm (blue diamonds) in z, while a low-resolution dataset at 80 nm spacing in z covered the entire volume (black dots). In C556 **(a5)**, both high- and low-resolution datasets covered the same region in z, had the same distance between slices, and no slices were discarded for any – therefore are all shown as black dots. **(b)** Warping datasets of different modalities to a common space using manually seeded landmarks. Details of the same region in both modalities and a group of local landmarks in context are shown for fixed 2-photon microscopy datasets warped into lab X-ray CT **(b1-3**, orange landmarks), lab X-ray CT into synchrotron X-ray CT **(b4-b5**, green landmarks) and in vivo 2-photon anatomical dataset onto serial block-face EM **(b6-b7**, blue landmarks). The landmarks are shown in **(b8-9)** in the context of the datasets involved, with the field of view of the 2-photon datasets outlined in blue and the tissue boundaries in X-ray or EM datasets outlined in grey.

**Supp. Fig. 6.**
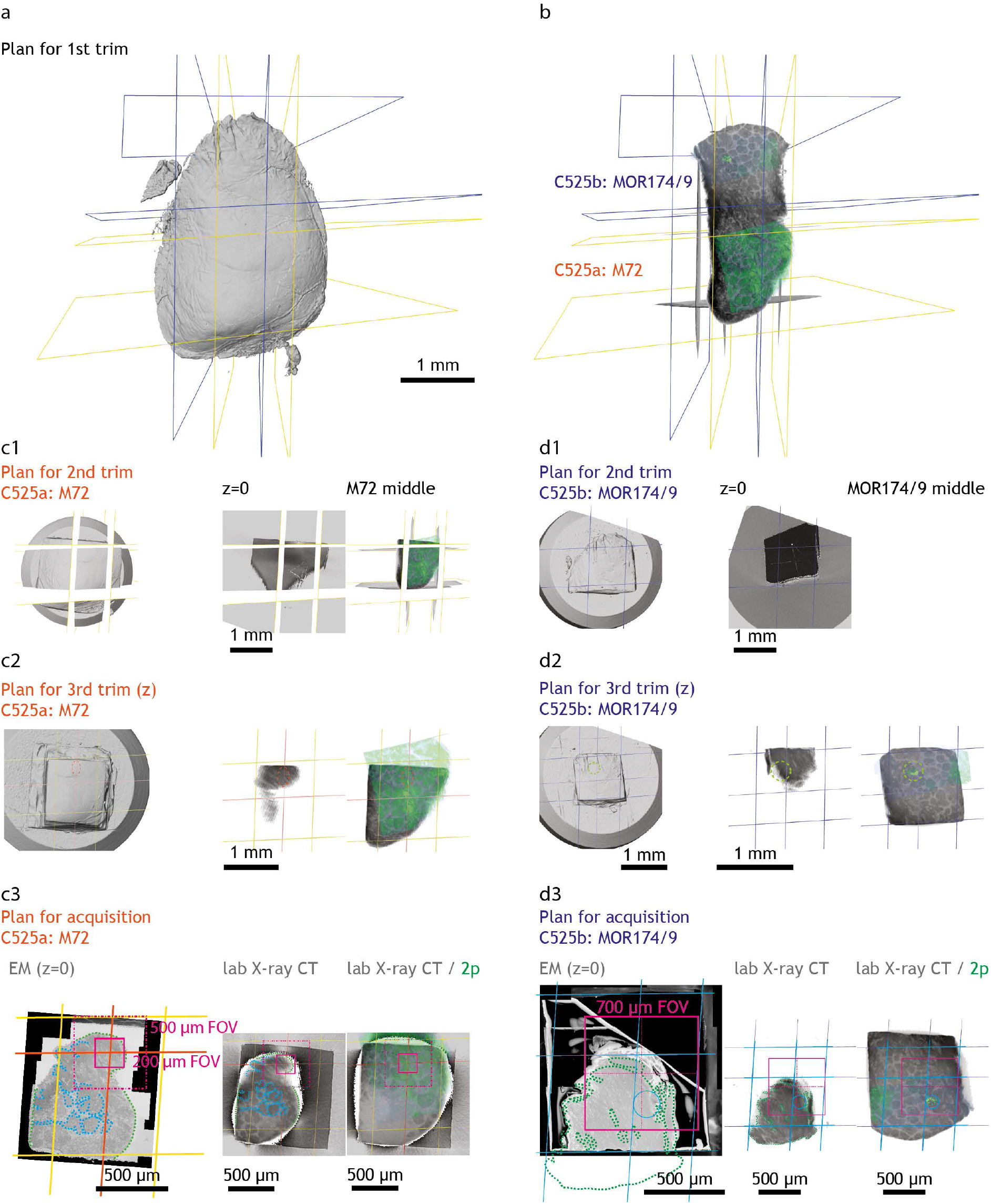
Specimen trimming into a geometry compatible with serial block-face EM. **(a)** Volume reconstruction of the X-ray CT dataset of the stained slab. A virtual section is shown in **(b)**, overlaid with warped 2-photon datasets taken of the fixed tissue, which carry information on the location of two genetically-tagged glomeruli, M72 and MOR174/9. Regions that contained both circuits were virtually resliced with yellow and blue planes, respectively. These images were then used to guide the mechanical trimming of the specimen, which was again scanned with lab X-ray CT, the dataset warped into the common framework, and iteratively trimmed until the sample was <1 mm^3^ in size for both M72 **(c1)** and MOR174/9 **(d1)** specimens. The block-face was then carefully shaved off (c2, d2) rendering a specimen with enough exposed tissue in the block-face to accurately predict the location of the ROI underneath (c3, d3). This last block-face would enable to configure the field of view of the datasets to acquire under the SBEM.

**Supp. Fig. 7.**
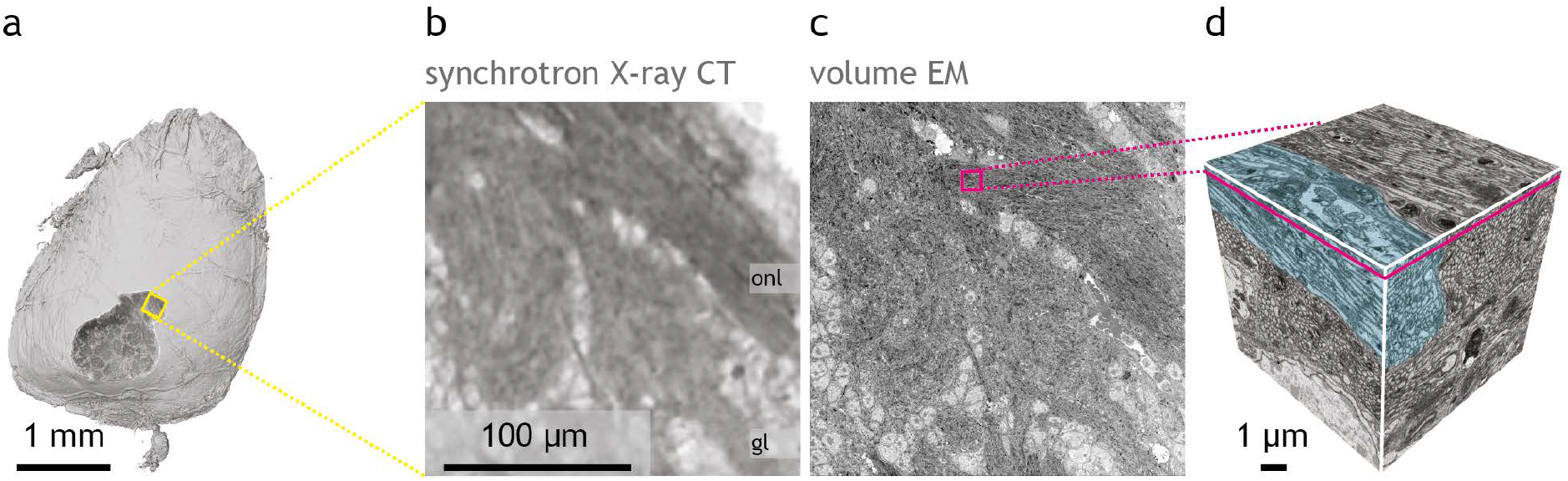
Ultrastructure is preserved after synchrotron X-ray CT. Volume of the whole specimen imaged by synchrotron X-ray CT **(a)** virtually sliced and a detail shown in **(b)** displaying olfactory nerve layer and glomeruli. **(c)** Same scene as resolved by serial block-face EM and volume shown in **(d)** displaying a bundle of axons of olfactory sensory neurons. *onl*, olfactory nerve layer; *gl*, glomerular layer.

**Supp. Fig. 8.**
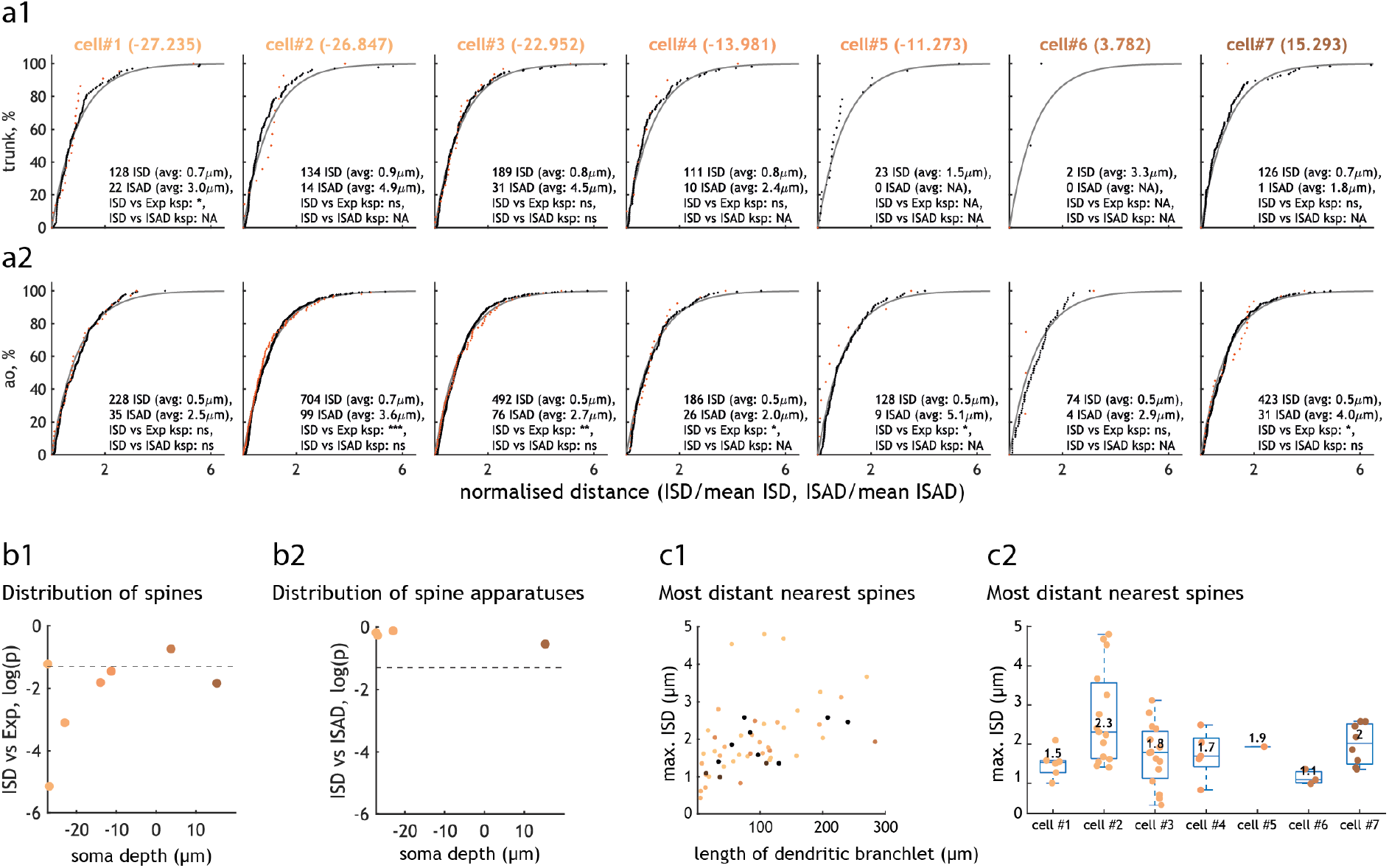
Distribution of spines and spine apparatus in the apical dendrites of CA1 pyramidal neurons. **(a)** Cumulative distribution of the inter-spine distance (ISD, black dots) and inter-spine apparatus distance (ISAD, red dots) on trunk **(a1)** and apical oblique **(a2)** dendrites of the seven CA1 pyramidal neurons traced and analysed at spine level (soma depth colour-coded in cell name as in **Fig. 5b**). For direct comparison, both distributions are normalised by their mean value (printed inside each plot). ISD distribution is also compared to an exponential fit of λ =1 shown in grey. An exponential distribution of ISD values would be expected if spines located randomly along the dendritic shaft. Similarly, ISAD distribution would follow ISD distribution if spine apparatus were located in a random subset of spines. Statistical significance of ISD versus exponential null and ISD versus ISAD are calculated using one-sample and two-sample Kolmogorov-Smirnov tests respectively, if there are more than 30 data points in each distribution (otherwise noted as NA). The p values obtained from these tests (“ksp”) are printed inside each plot. **(b)** Probability that the distribution of spines **(b1)** and spine apparatus **(b2)** in apical oblique dendrites is random. Values depict, for each cell with its own soma depth, the p value from one-sample Kolmogorov-Smirnov test between inter-spine distance distribution and a fitted exponential distribution **(b1)** and for cells that have more than 30 spine apparatus in their apical oblique dendrites the p value from two-sample Kolmogorov-Smirnov test between inter-spine and inter-spine apparatus distributions **(b2)**. (* p<0.05, ** p<0.005, *** p<0.0005, ns nonsignificant). **(c)** Maximal inter-spine distance, as a function of the length of each dendritic branchlet analysed **(c1)** and grouped per cell **(c2)**. For each cell, the median is displayed in the box plot. Each dendritic branchlet is represented by a dot, dots coloured according to the soma depth of the cell. Note there is an upper bound on inter-spine distances, with no spines found further apart than 2 μm in most dendrites of all cells analysed. Large ISDs are therefore less frequent than expected by chance.

**Supp. Fig. 9.**
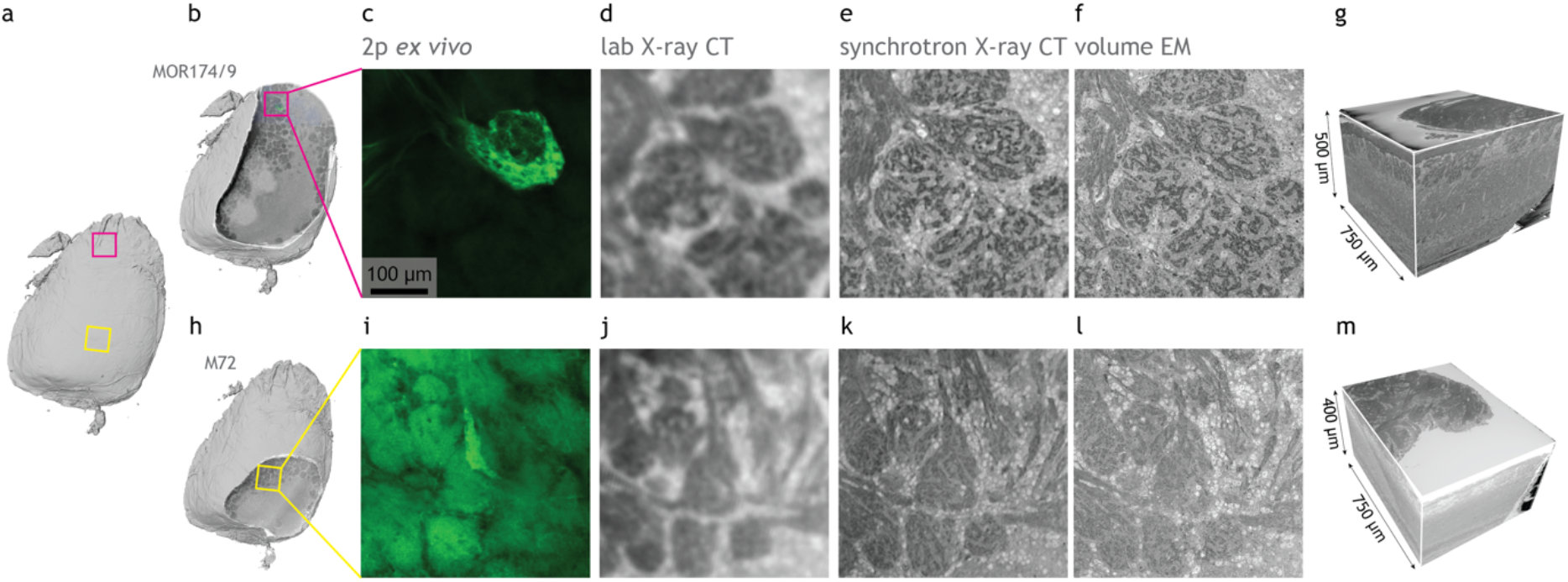
Multiple serial block-face EM datasets obtained from the same specimen and linked to the same 3D environment. **(a)** Reconstruction of an olfactory bulb slab, virtually sliced in **(b,h)** displaying the regions of interest containing the MOR174/9 and M72 glomeruli in magenta and yellow, respectively. **(c-f)** Multimodal correlated image data of the same region, displaying 2-photon dataset of the fixed specimen **(c)**, lab X-ray CT **(d)**, synchrotron X-ray CT **(e)** and serial block-face EM **(f)** of the MOR174/9 region of interest delineated in **(b)**. **(g)** Reconstruction of the SBEM dataset containing the MOR174/9 glomerular column. **(i-l)** Multimodal correlated image data of the same sample but for the M72 region of interest delineated in (h). Modalities as in (c-f). (m) Reconstruction of the SBEM dataset containing the M72 glomerular column.

**Supp. Fig. 10.**
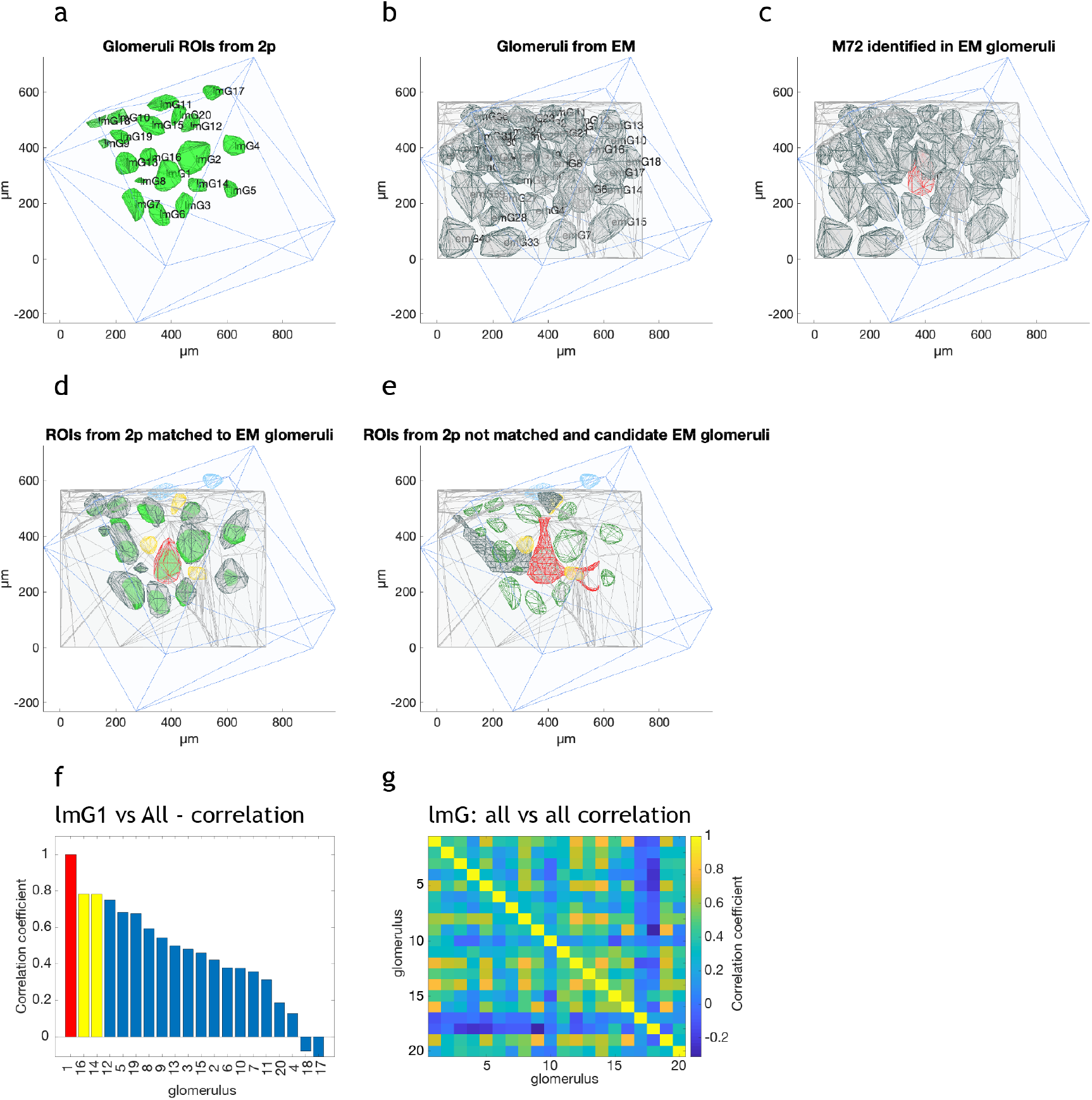
Tracking glomerular regions of interest between 2-photon and serial block-face EM. **(a)** Glomerular ROIs segmented in the two-photon dataset. ROIs shown in green, with their respective 2-photon identifier tags overlapped. The field of view covered by the 2-photon dataset is shown in blue. **(b)** Glomerular ROIs segmented in the EM dataset. ROIs shown in grey, with their EM identifier tags overlapped. The field of view covered by the EM dataset is shown in pale grey. **(c)** The glomerulus M72 is highlighted in red. **(d)** Matched glomerular ROIs between 2-photon and EM. Only matched EM ROIs are shown, along with matched 2-photon (green), unmatched 2-photon (yellow) and 2-photon ROIs that fall outside the EM field of view (blue). **(e)** Detailed contours of EM ROI candidates for the unmatched 2-photon ROIs. Two unmatched ROIs fall very close to axon bundles associated with the M72 glomerulus, which also contains a matched ROI. **(f)** Cross-correlation between the activity profile recorded in the M72 ROI and in all other ROIs. The columns of the two nearby unmatched ROIs are highlighted in yellow. **(g)** Cross-correlation of the activity profile recorded across all glomeruli.

**Supp. Fig. 11.**
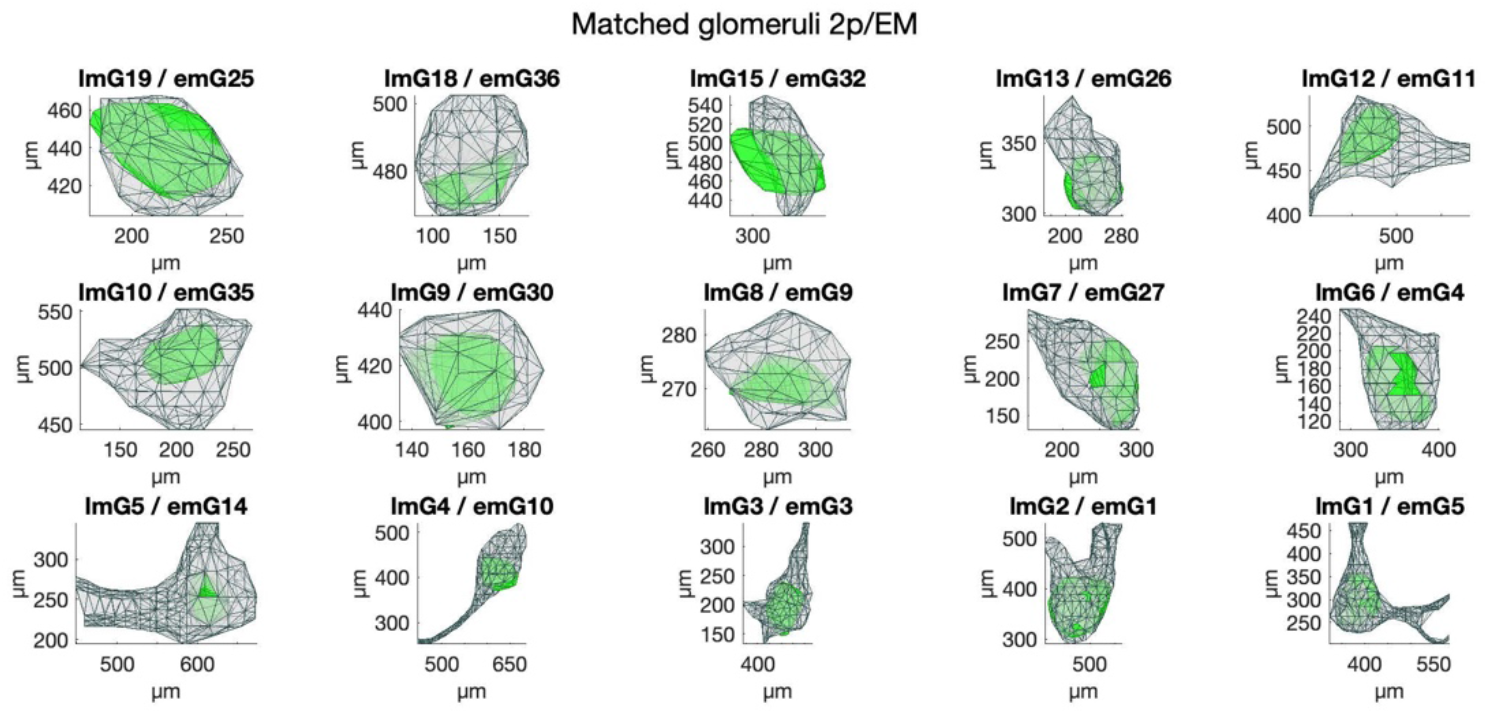
Matched glomerular regions of interest between 2-photon and serial block-face EM. Individual plots for all matched glomerular ROI pairs 2-photon (green) / EM (grey).

